# Rethinking the nature of intraspecific variability and its consequences on species coexistence

**DOI:** 10.1101/2022.03.16.484259

**Authors:** Camille Girard-Tercieux, Isabelle Maréchaux, Adam T. Clark, James S. Clark, Benoît Courbaud, Claire Fortunel, Joannès Guillemot, Georges Künstler, Guerric le Maire, Raphaël Pélissier, Nadja Rüger, Ghislain Vieilledent

## Abstract

**Context:** Intraspecific variability (IV) has been proposed to explain species coexistence in diverse communities. Assuming, sometimes implicitly, that conspecific individuals can perform differently in the same environment and that IV blurs species differences, previous studies have found contrasting results regarding the effect of IV on species coexistence.

**Objective:** We aim at showing that the large IV observed in data does not mean that conspecific individuals are necessarily different in their response to the environment and that the role of high-dimensional environmental variation in determining IV has been largely underestimated in forest plant communities.

**Methods and Results:** We first used a simulation experiment where an individual attribute is derived from a high-dimensional model, representing “perfect knowledge” of individual response to the environment, to illustrate how a large observed IV can result from “imperfect knowledge” of the environment. Second, using growth data from clonal *Eucalyptus* plantations in Brazil, we estimated a major contribution of the environment in determining individual growth. Third, using tree growth data from long-term tropical forest inventories in French Guiana, Panama and India, we showed that tree growth in tropical forests is structured spatially and that despite a large observed IV at the population level, conspecific individuals perform more similarly locally than compared with heterospecific individuals.

**Synthesis:** As the number of environmental dimensions that are typically quantified is generally much lower than the actual number of environmental dimensions influencing individual attributes, a great part of observed IV might be misinterpreted as random variation across individuals when in fact it is environmentally-driven. This mis-representation has important consequences for inference about community dynamics. We emphasize that observed IV does not necessarily impact species coexistence *per se* but can reveal species response to high-dimensional environment, which is consistent with niche theory and the observation of the many differences between species in nature.

## Introduction

Ecological communities are characterized by numerous coexisting species, for instance in grasslands, coral reefs or tropical forests. Understanding how these species stably coexist while competing for the same basic resources, *viz*. light, water, and nutrients (Baraloto et al. 2010), is a longest-standing question in ecology (Gause 1934, Hutchinson 1961, Levine et al. 2017). Although numerous mechanisms have been suggested to contribute to species coexistence (Janzen 1970, Connell 1971, Chesson 2000, Hubbell 2001, Wright 2002, Levine and HilleRisLambers 2009), it is unclear when and to what extent they explain the high species diversity observed in nature (Clark 2010). This is especially true in forests, where tree species coexist while seemingly requiring similar resources in the same location. Astonishingly, a hectare of tropical forest can harbor more than 900 plant species of a diversity of forms and functions (Wilson et al. 2012).

Many theoretical mechanisms that might explain tree species coexistence typically follow the assumption that all conspecific individuals are identical. However, intraspecific variability (IV) in traits, demographic rates or any proxy of performance, henceforth denoted as “attributes”, can alter community structure and dynamics (Bolnick et al. 2011). Indeed, large IV has been observed across a number of attributes in plant communities(Albert et al. 2012, Violle et al. 2012). For instance, Siefert et al. 2015 estimated that IV accounted for 25% of the variability in functional traits within plant communities on average, and this proportion was even estimated at 44% in a tropical forest (Poorter et al. 2018). Likewise, IV in growth rates for trees of standardized size, local crowding and terrain slope has been found to account for up to 58% of total growth variability in a tropical forest stand (Le Bec et al. 2015).

IV, as a pathway for coexistence, has so far not shared the same attention as other mechanisms. This is in part because modeling studies that have explored the effect of IV on species coexistence have yielded contrasting results (Stump et al. 2021). In most theoretical analyses, variability in attributes among conspecific individuals has been included through independent random draws (Lichstein et al. 2007, Hart et al. 2016, Barabás and D’Andrea 2016, Crawford et al. 2019 but see Purves and Vanderwel 2014, Banitz 2019). Similarly, empirical studies typically summarize IV as a variance around species mean attributes (Jung et al. 2010, Albert et al. 2010, Siefert et al. 2015, Poorter et al. 2018). With this representation, IV can increase species niche overlap and blur species differences, sometimes slowing down competitive exclusion in models of community dynamics (Vieilledent et al. 2010, Crawford et al. 2019). However, in some other models, non-linear responses can make such IV beneficial to the superior competitors (*i*.*e*. the most competitive individuals of the more competitive species), thus accelerating competitive exclusion (*e*.*g*. Courbaud et al. 2012, Hart et al. 2016). Alternatively, in particular spatial configurations, more precisely when IV is greater in species preferred habitats, it has been shown to foster species coexistence (Uriarte and Menge 2018). Stump et al. (2021) have proposed to reconcile these contrasting results by distinguishing the effect of IV on niche traits (which control individual performance response to environmental conditions) vs. hierarchical traits (which control individual performance independently from environmental conditions). They demonstrated with different simulation models of community dynamics that IV in traits can alter stabilizing mechanisms and fitness differences in a complex way which depends upon the nature of the traits (niche vs. hierarchical) and their response curve, and thus promote or not species coexistence. In all the above examples however, IV, since simulated through independent random draws around species mean attributes, would be caused by differences among individuals that are fully independent of the environment: differences among individuals would remain unchanged even when experiencing exactly the same environmental conditions. Importantly, such simulated IV thus leads to a variation among conspecific individuals that is completely unstructured in space and time. New appreciation of fine-scale environmental heterogeneity and structure as well as species differences in their response to the environment, however, may suggest that this assumption of unstructured IV is rarely met.

Novel remote sensing tools such as high-spatial and -temporal resolution airborne LiDAR scans (Tymen et al. 2017, Cushman et al. 2022), intensive soil samplings and metabarcoding (Zinger et al. 2019), and more generally studies on the microclimate (Zellweger et al. 2019) and microhabitats (Baraloto and Couteron 2010) have indeed evidenced strong environmental variation operating at fine scales (*e*.*g*. cm to meter scales) in many dimensions (Fig. 1). These environmental dimensions can be resources for which species compete (*e*.*g*. light, water, nutrients) but also all other components that shape the environment locally in space and time (*e*.*g*. temperature, wind, elevation, slope, soil texture, soil microorganisms *etc*.). In parallel, naturalists and taxonomists have long documented species differences in many aspects of their morphology and life history (Fig. 2). Such differences between species have then been specified and quantified through traits that drive each species response to the environment (species functional traits, Mcgill et al. 2006, Westoby and Wright 2006). Similar to the environment that presents highly-dimensional variation at local scales, these functional species differences spread along many dimensions within communities (Hutchinson 1957, 1959, Baraloto et al. 2010, Kraft et al. 2015, Rüger et al. 2018, Maréchaux et al. 2020, Vleminckx et al. 2021).

**Figure 1:**
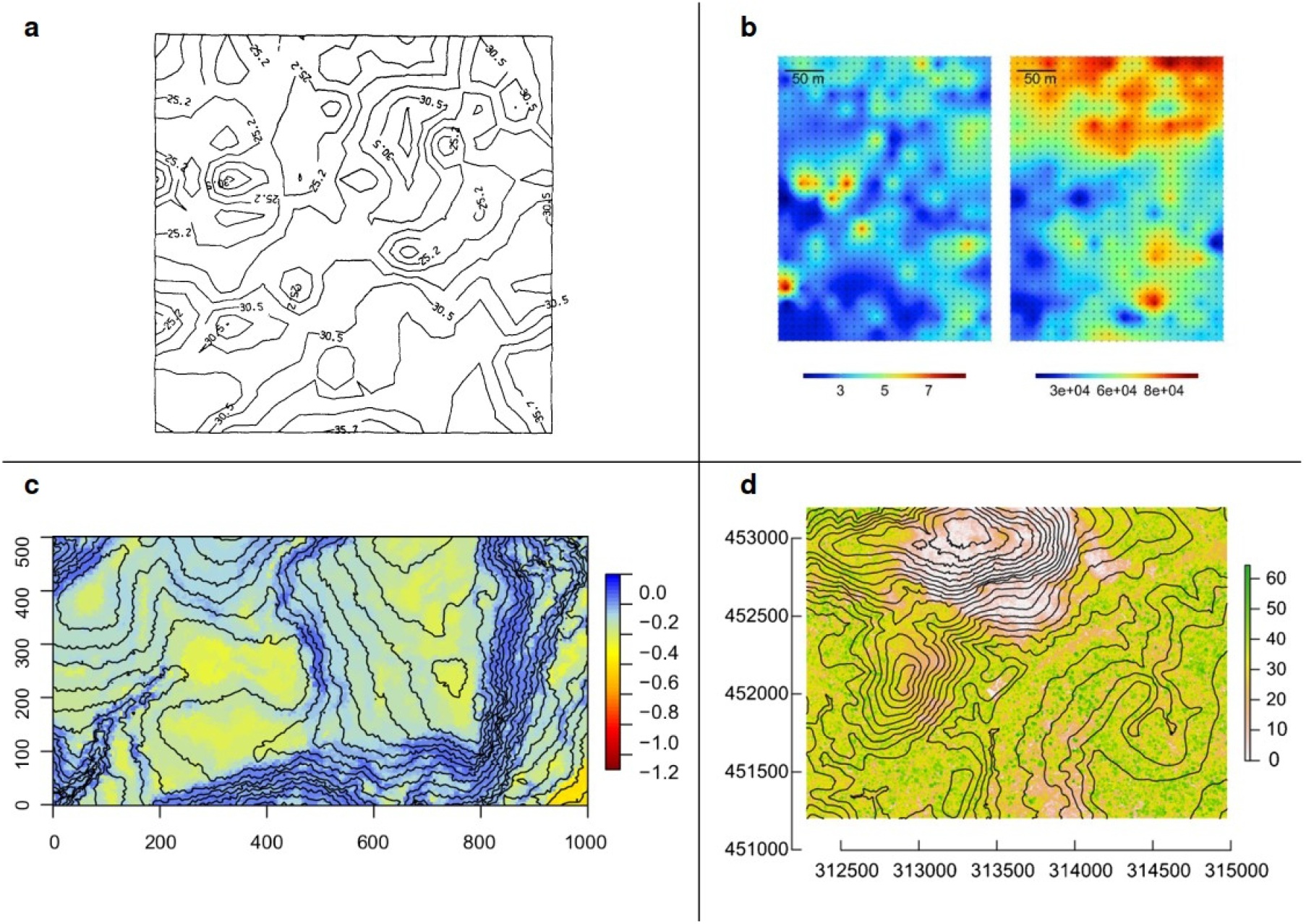
High environmental variability at a small spatial scale. **(a)** Soil nitrogen content in a 12×12 m plot at Cedar Creek in g.kg^−1^, (USA), Tilman 1982; **(b)** Carbon in % (left) and aluminum in ppm (right) soil content in a 12-ha (250×500 m) plot at The Nouragues (French Guiana), Zinger et al. 2019; **(c)** Soil water content during mid-dry season of a regular year in MPa in a 50-ha (1000×500 m) forest plot at Barro Colorado Island (Panama), Kupers et al. 2019. Coordinates in m.; **(d)** Canopy height in m and topography (10 m spaced elevation lines) in a 50-ha (2500×2000 m) area at the Nouragues Research Field Station, Tymen et al. 2017. Coordinates in m (UTM 22N).

**Figure 2:**
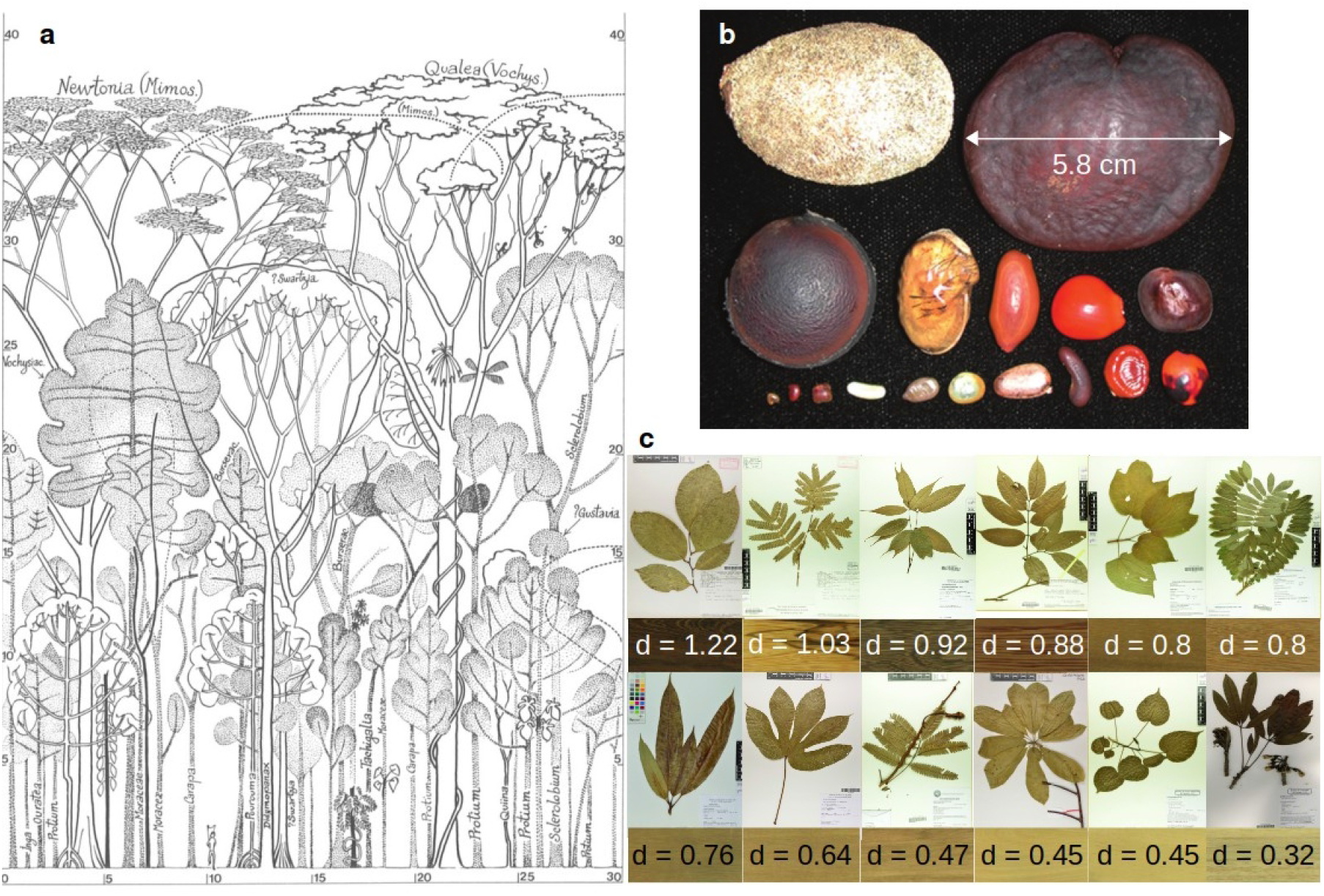
Morphological diversity of tree species illustrating strong differences between species. **(a)** Diversity of tree species architecture and height in a tropical forest (Hallé et al. 1978). Coordinates are in m.; **(b)** Diversity of seed size and shape from 17 tree species of the Fabaceae family in the Peruvian Amazon (Muller-Landau 2003); **(c)** Diversity of leaf size and shape (herbarium of Cayenne, Gonzalez et al. 2021) and of wood aspect (reflecting wood characteristics) and density (Normand et al. 2017) for 12 tree species in French Guiana. Species from top left to bottom right are *Bocoa prouacensis, Zygia racemosa, Vouacapoua americana, Eperua falcata, Bagassa guianensis, Hymenolobium excelsum, Mangifera indica, Sterculia pruriens, Parkia nitida, Couroupita guianensis, Hura crepitans*, and *Ceiba pentandra*. Black bars next to herbarium samples indicate the scale (10 cm).

In this paper, we explore the potential that the role of environmental variation in shaping *observed* IV has been largely underestimated with important consequences on our understanding of the effect of *observed* IV on community dynamics. Indeed, a great part of *observed* IV might emerge from species responses to a high-dimensional environment (Fig. 3): observed differences among individuals of the same species can be caused by the (often poorly quantified) differences in the micro-environment they experience. If so, variation among conspecific individuals would be structured in space and time, and not necessarily by genetic variation. More specifically, we present insights from a simulation experiment, experimental data, and tropical forest inventory data in order to examine three hypotheses (Fig. 4): (i) the large IV observed in natural communities can emerge from heterogeneity in multiple unobserved environmental dimensions; (ii) because environmental variation is structured in space and time, IV is likely to be similarly structured as well, suggesting that it is not appropriate to represent IV as a purely random noise in models; and (iii) since a large observed IV does not necessarily imply that conspecific individuals substantially differ in their fundamental niche, conspecific individuals may still respond more similarly to environment than heterospecific individuals. We therefore call for a reconsideration of the nature and structure of observed IV, which could shed new light on the coexistence conundrum. While we acknowledge the existence of genetically-based individual variations, and that plasticity has a genetic basis (Nicotra et al. 2010, Westerband et al. 2021), we suggest that a substantial part of *observed* IV might result from the higher dimensionality of the species niche than typically observed. Species differences along these many dimensions can lead to multiple local inversions of species hierarchy in an environment varying in space and time, thereby allowing the stable coexistence of numerous species.

**Figure 3:**
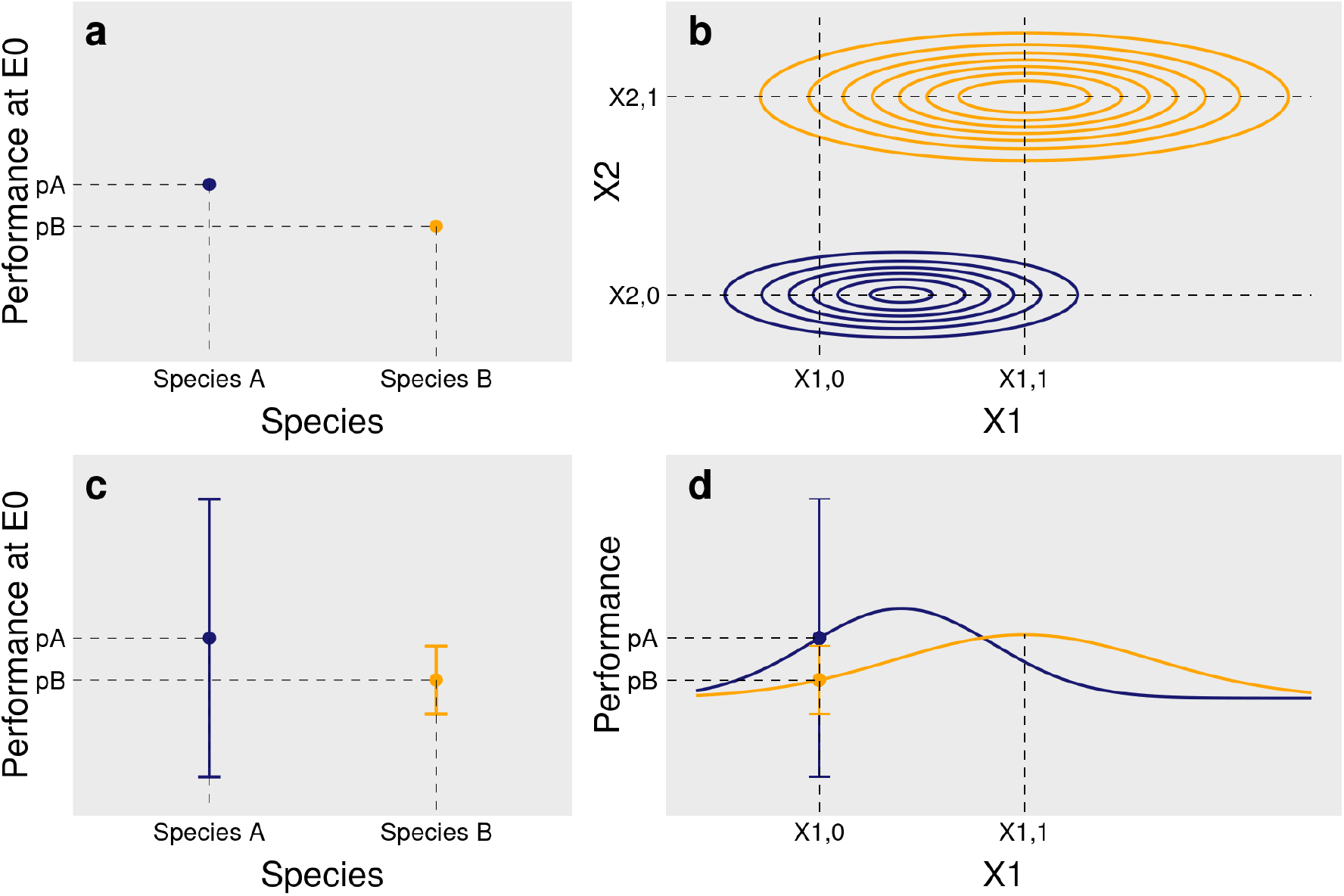
Reinterpreting observed intraspecific variability (IV): from niche widening to niche projection into a high-dimensional environment. In **(a)**, within a given environment E0 defined along an environmental axis X1 (E0=E(X1,0)), conspecific individuals are identical and have the same performance pA and pB, for species A (blue) and species B (orange). Species A outcompetes species B in E0. Actual measured differences among conspecific individuals, shown in **(c)**, can be interpreted in different ways. First, as conspecific individuals exhibit contrasting attributes in E0, they become more different. This can result in some heterospecific individuals having similar performances: IV would blur species differences. Alternatively, IV measured in E0 results from the variation of unobserved environmental variables (E0=E(X1,0, X2); **(b)**). Contrasting performances among conspecific individuals in E0 do not result from intrinsic differences among them but from differences in the local environment they experience and that was poorly characterized, *i*.*e*. the number of observed dimensions is lower than the actual number of environmental dimensions. Similarly, although species niches present some overlap when projected on one dimension **(d)**, they do not overlap in the two-dimensional space **(b)**. Moreover, while species A outcompetes species B on average when X1=X1,0, the opposite occurs when X1=X1,1 **(d)**, leading to an inversion of species hierarchy between different environments. Similarly, while species A outcompetes species B in E(X1,0, X2,0), the opposite occurs in E(X1,0, X2,1). Although only two dimensions are shown, species respond to many environmental variables varying in space and time, multiplying the possibilities of niche segregation and hierarchy inversions between species, offering room for species coexistence in a variable high-dimensional environment. The code used to generate this figure is available online.

**Figure 4:**
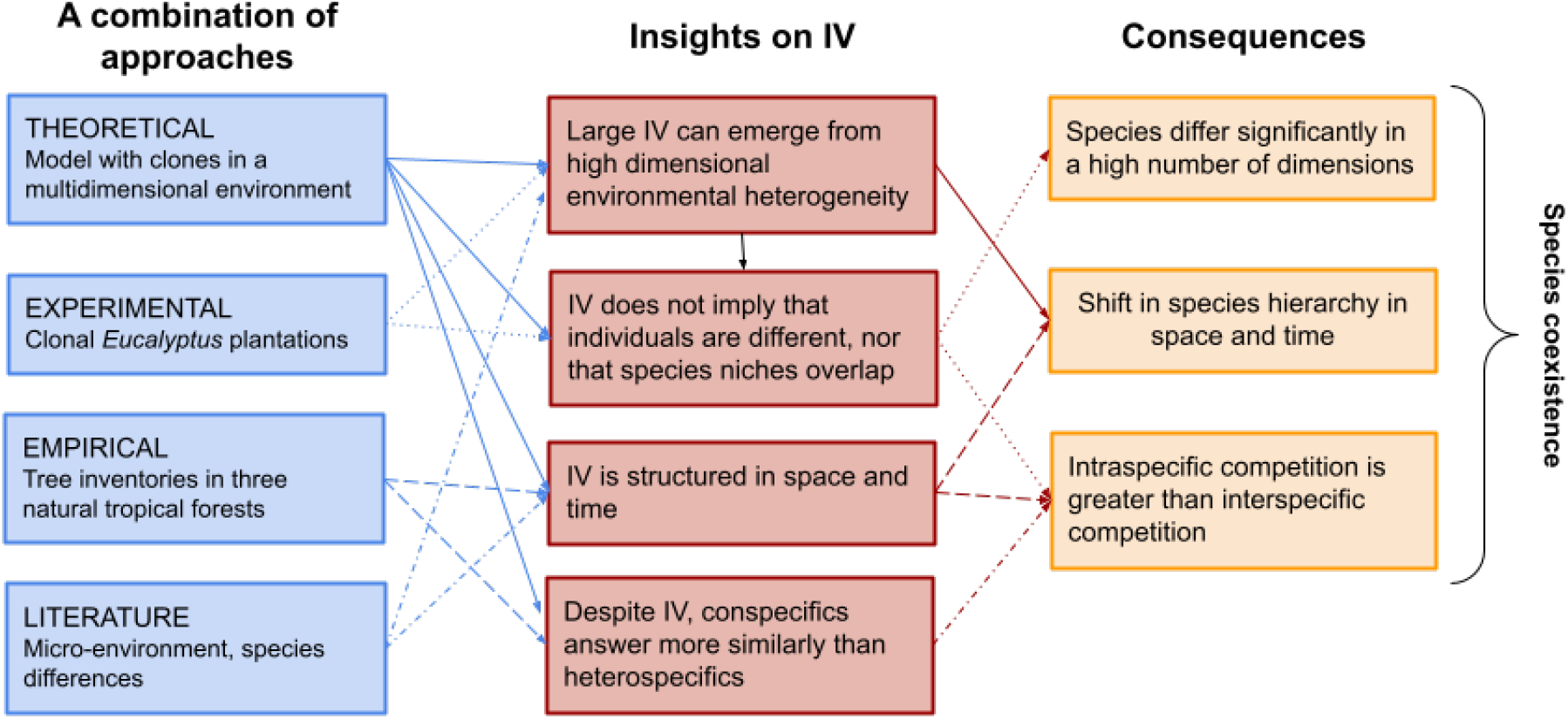
Multiple insights on the nature of IV and its consequences on individual and species differences. We used literature and data analyses of various nature to support the hypothesis that a large part of observed IV can result from multidimensional environmental variations that are spatially and temporally structured rather than by intrinsic and spatio-temporally unstructured differences between conspecific individuals, with radically different consequences on species coexistence.

## Theoretical illustration: unobserved environmental dimensions result in large observed IV

We first conducted a simulation experiment to illustrate how observed intraspecific variability, or “individual effects”, can result from variation in unobserved environmental variables (as suggested by (Clark et al. 2007). We generated simulated data of an individual attribute (here tree growth) depending on a certain number of environmental variables varying in space, and then analyzed the simulated data assuming that most of the environmental variables are actually unobserved, as it is typically the case in the field.

### A “perfect knowledge” simulation model

We considered a set of *J* species with *I* individuals each, distributed in a virtual landscape. The environment was assumed to be fully known and defined by *N* environmental variables, *X*_*1*_ to *X*_*N*_, that were each randomly and independently generated in the landscape, assuming spatial autocorrelation. Individual location was drawn randomly in a virtual landscape defined by a *C* × *C* square grid, each cell corresponding to a particular environment (Fig. 5a). Individuals were identical within species (same model parameters for all conspecific individuals), but different between species (different model parameters between heterospecific individuals).

**Figure 5:**
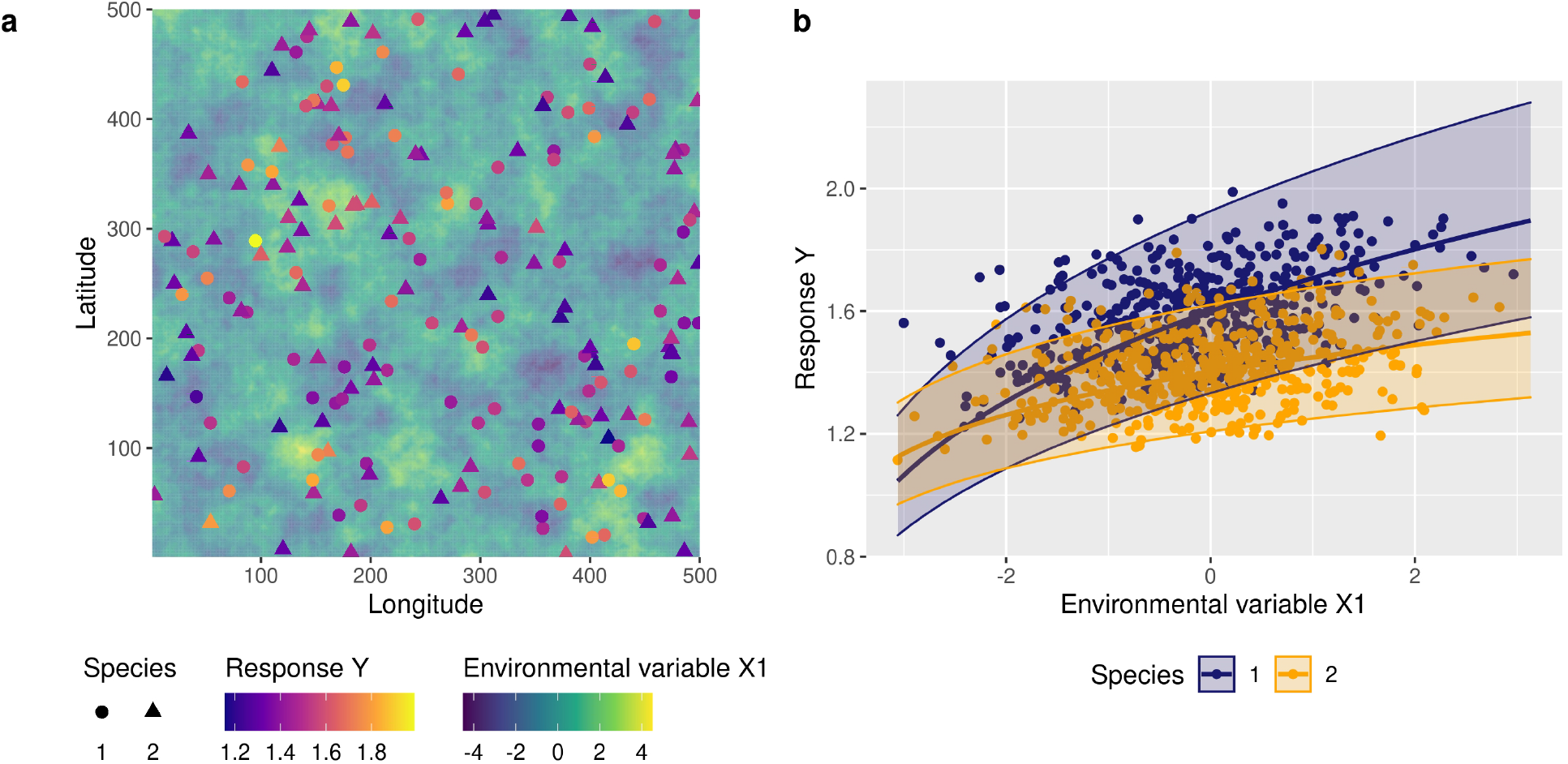
Observed intraspecific variability as a result of the imperfect characterization of the environment. A simulated response variable (Y, *e*.*g*. growth) is generated for individual clones of two species thriving in a high-dimensional environment. This response variable was first computed as a function of ten environmental variables (“perfect knowledge” model, Eq. I), but is then analyzed using a statistical model that accounts for the unique environmental variable that was assumed to be observed in the field (X_1_, *e*.*g*. light) and includes a random individual effect (“imperfect knowledge” model, Eq. II). The intraspecific variability estimated with these random individual effects is then due to the variation in space and time of the nine unobserved environmental variables. **(a)** Positions of a sample of I=600 individuals from J=2 species in a landscape defined by a square grid of C × C cells (C=500). The background color indicates the value of the observed environmental variable X_1_ on each cell at date t. The response Y of each individual, which depends on the environment within each cell (Eq. I), is also indicated by a color scale. **(b)** Response Y as a function of the observed environmental variable X_1_ for the two species. Points represent the data {Y_ijt_, X_1,ijt_}. Thick lines represent the predictive posterior means for the two species. The envelopes delimited by two thin lines represent the 95% credible intervals of the predictive posterior marginalized over individuals (taking into account 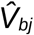). The envelopes thus represent the intraspecific variability which is due to the N-1 unobserved environmental variables.

We considered the following “perfect knowledge” mathematical model, which depicts the exact attribute *Y*_*ijt*_ (*e*.*g*., growth) of an individual *i* of species *j* given its environment at time *t* (Eq. I, Appendix 1).

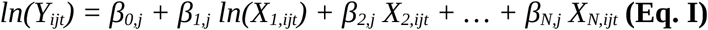

In this model, *β*_*j*_ = [*β*_*0,j*_, …, *β*_*N,j*_] is the vector of parameters defining the response of individuals of species *j* to the environment. Because conspecific individuals respond similarly to environmental variables, variation in *Y*_*ijt*_ among them is only due to differences in the environment where and when each individual is growing. Using this model, we computed the attribute *Y* of the *I*×*J* individuals at *T* dates, assuming that values for some of the environmental variables changed between dates, and thus obtained a simulated dataset {*Y*_*ijt*_, *X*_*1,ijt*_, …, *X*_*N,ijt*_} with *N*=10, *I* = 300, *J* = 2, *C* = 500 and *T* = 2.

### An “imperfect knowledge” statistical model

Second, we considered an “imperfect knowledge” statistical model for which we assumed that only one explanatory variable *X*_*1*_ (*e*.*g*., light) in the above simulated dataset has been measured among all the environmental drivers that actually determine response variable *Y* (Eq. II, Appendix 1). This model represents the ecologist’s imperfect understanding of attribute *Y*. The model includes a species fixed effects on the intercept and on the slope (*β’*_*0,j*_ and *β’*_*1,j*_) and a random individual effect b_0,i_ on the intercept, *b*_*0,i*_ ∼ N(0, *V*_*bj*_), where *V*_*bj*_ is the intraspecific variance for species *j*. We estimated the model parameters based on the simulated dataset introduced above but considering only the first explanatory variable {*Y*_*ijt*_, *X*_*1,ijt*_}, the remaining “unknown” environmental effects being contained in the model residuals, *ε*_*ijt*_.

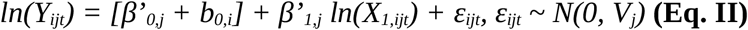

### Apparent niche overlap and observed intraspecific variability as a result of unobserved environmental variables

Despite the fact that conspecific individuals were identical and species responses to environment were different, the variance estimates 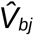 for individual random effects of species *j* were large, and species responses to the environment overlapped (Fig. 5b). This is due to the contribution of the unmeasured variables {*X*_*2,ijt*_, …, *X*_*N,ijt*_} in determining the variation of *Y* across individuals.

Since it is driven by spatially autocorrelated variables (Eq. I), the response *Y* was spatially autocorrelated across conspecific individuals (Fig. 6). This means that two neighboring conspecific individuals have more similar attribute *Y* than two distant conspecific individuals. Additionally, the variance of *Y* was lower within than between species: conspecific individuals responded more similarly to the environment than heterospecific individuals did (Fig. 6).

**Figure 6:**
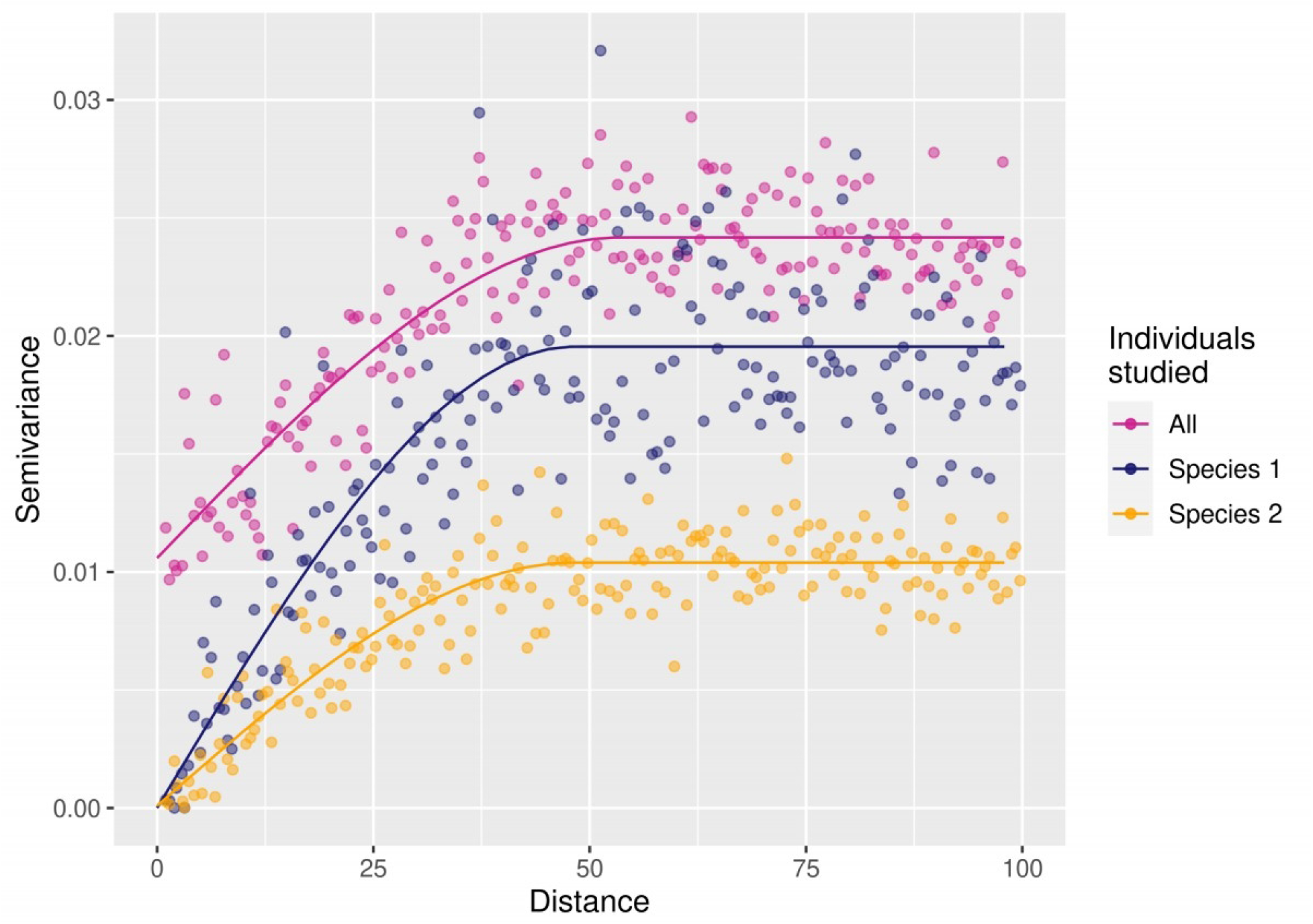
Spatial autocorrelation of attribute Y across individuals within and between species (J=2) in a simulation experiment. This semivariogram represents the semivariance of the individual mean attribute Y as a function of the distance between individuals. The increasing curves evidence spatial autocorrelation in Y (similar results using Moran’s I test). The semivariance of all individuals taken together (purple curve) is higher than the semivariance of conspecific individuals for the two species (orange and blue curves), which means that intraspecific variability is lower than interspecific variability.

With this simulation experiment, we simply illustrated that: (i) a high IV can emerge from unobserved environmental dimensions exclusively, (ii) the spatial structure of IV follows the spatial structure of the underlying environmental variables, and (iii) IV does not blur differences between species (Fig. 6) despite apparent niche overlap (Fig 5b).

## Experimental insights: large observed intraspecific variability in a clonal tree plantation

We then moved from a theoretical to an experimental approach using census data from clonal *Eucalyptus* plantations, where genetic variability among individuals growing within a single same site is controlled. We explore the partitioning of IV between intrinsic (genotypes) and extrinsic sources, which is often infeasible in natural settings, to demonstrate that substantial observed IV can indeed emerge from genetically identical individuals in the field, even when persisting in an apparently homogeneous environment.

### An extreme case of controlled genetic and environmental variation

The EUCFLUX experiment (São Paulo state, Brazil) is a clonal trial with a replicated, statistically-sound design (le Maire et al. 2019). It includes 14 genotypes of 5 different *Eucalyptus* species or hybrids of various origins. Each genotype is planted in plots of 100 trees, at a density of 1666 trees per hectare, and replicated spatially in 10 blocks (Fig. 7). The experimental set-up was designed to minimize the variation in environmental factors among blocks, which were separated by less than 1.5 km within a homogeneous 200-ha stand showing small variation in soil properties. Tree diameter at breast height (*D*) has been measured over 5 complete censuses, spanning 6 years, age at which such plantation is generally harvested (see le Maire et al. 2019 and Appendix 2 for further details on this experimental set-up).

**Figure 7:**
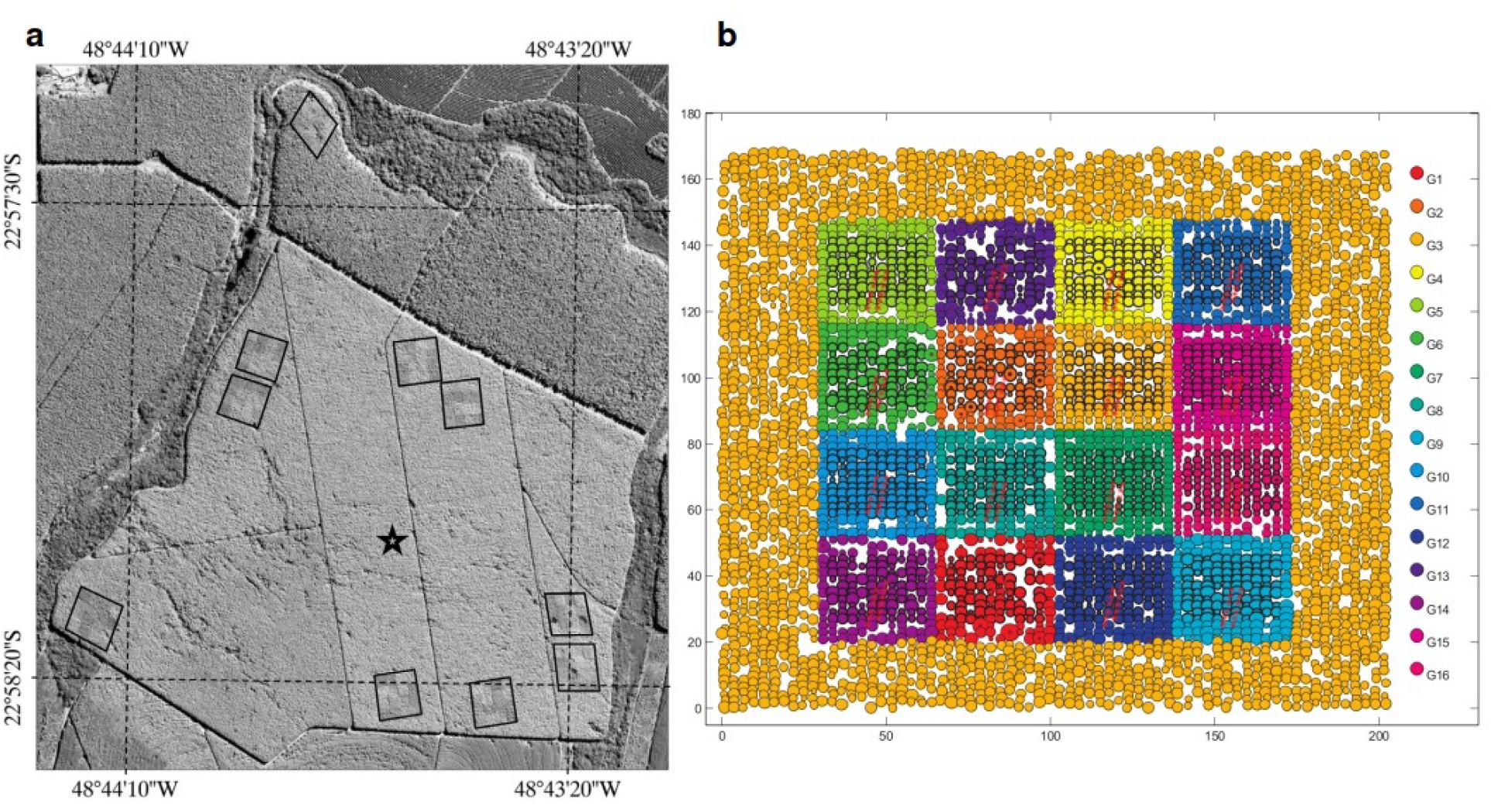
Experimental setup of the EUCFLUX experiment. The ten blocks **(a)** and the organization of the 16 genotypes within a block **(b)**. In our analyses, two genotypes were discarded because they were obtained from seeds and not clones and therefore included some genetic variability. A more complete figure legend can be found in le Maire et al. 2019.

### A partitioning of observed variance among individual tree growth

We computed annual diameter growth (*G*) in mm/yr^-1^ for each tree as well as a competition index (*C*) as the sum of the basal area of the eight direct neighbors of each tree. The dataset included 64,125 growth estimates corresponding to 13,531 trees in total. To quantify the relative importance of the different sources of growth variability, we used a statistical hierarchical growth model (Eq. III) including an intercept (*β*_*0*_), fixed effects of the log-transformed diameter (*β*_*1*_) and competition index (*β*_*2*_), and random effects on the intercept of the block (*b*_*0,b*_, with *b*_*0,b*_ *∼ N(0, V*_*b*_)), the genotype (*b*_*0,g*_, with *b*_*0,g*_ *∼ N(0, V*_*g*_)), the census date (*b*_*0,t*_, with *b*_*0,t*_ *∼ N(0, V*_*t*_)), and the individual (*b*_*0,i*_, with *b*_*0,i*_ *∼ N(0, V*_*i*_)). All the data were log-transformed and scaled, and a constant of 1 mm was added to all growth values to avoid undefined logarithms.

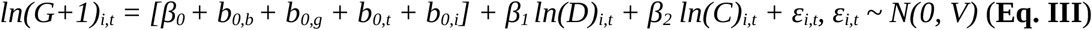

We used conjugated priors with inverse-gamma distributions (with shape and scale parameters=10^−3^) for variance parameters, and normal distributions (with mean=0 and variance=1) for mean parameters. The estimation of model parameters was done using a Bayesian approach using Stan software with the brms R package (Bürkner 2017, 2018). We made 10,000 iterations for each MCMC with a burn-in period of 5,000 steps and a thinning rate of one fifth. We obtained 1,000 estimations per parameter and examined the trace plots to check convergence of the MCMC chains.

We then examined the proportion of the model residual variance (variation of the response variable that is not explained by the covariates) related to each random effect in order to partition the block, genotype, date and individual variances.

### Variation among individuals is not explained by genotype

While minor variability was associated with blocks (Table 1), confirming that they are broadly homogeneous by design, the variability associated with extrinsic temporal factors was predominant (Table 1). It reveals that the competition index (*C*) used in the analysis to encapsulate the effect of progressive canopy closure does not fully encompass all temporal effects.

**Table 1:**
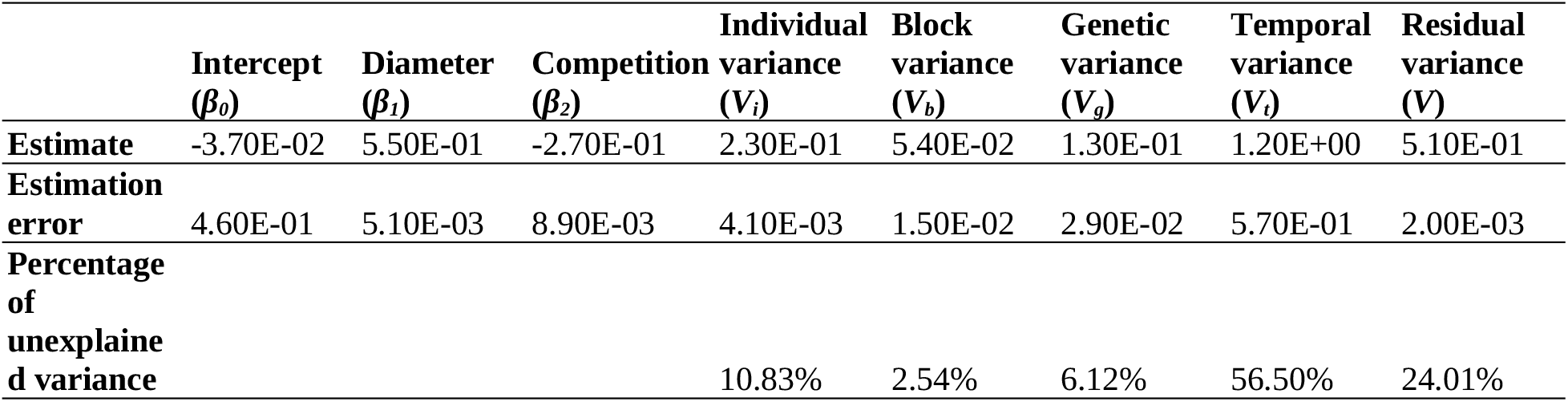
Mean posteriors of the *Eucalyptus* model and their estimation errors and residual variance partitioning among the different random effects.

Importantly, the variability between individuals was almost twice as high as the variability between genotypes (Table 1). Hence, even in such an extremely conservative case, where environmental variation in space is minimized and genotypic variability controlled, a large part of measured IV cannot be explained by purely-genetic differences among individuals that would remain independent of the environment as an IV simulated through independent random draws would be. This suggests an underestimated role of environmental micro-heterogeneity in shaping variation among individuals, for instance inevitable spatial variation of biotic and abiotic variables (soil microbiome, pathogens, soil structure and water content, light *etc*.) at fine scales (e.g. cm- to m-scale, hence impacting tree-scale environment, Baraloto and Couteron 2010, Fig. 1) as well as potential early manipulations of the young plant, the way it was planted, *etc*.

## Empirical insights: observed intraspecific variability is high and spatially structured and does not blur species differences in tropical forests

To test some of our hypotheses in natural communities, we then used data from three long-term tree inventories in tropical forests, from Amazonia (Paracou, French Guiana; Gourlet-Fleury et al. 2004), Central America (Barro Colorado Island, Panama; Losos and Leigh 2004) and South-East Asia (Uppangala, India; Pélissier et al. 2011). More specifically, we inferred observed IV, tested if individual growth showed local spatial autocorrelation, *i*.*e*. was structured in space, and if conspecific individual growth was more similar than heterospecific individual growth locally. These three sites encompass contrasting climatic conditions (rainfall ranging from 2,600 in BCI to 5,100 mm.y^-1^ in Uppangala), disturbance regimes (incl. various logging experiments in Paracou) and topography (from gentle in BCI to mountainous in Uppangala), making them representative of the global tropical forests. The data from these tropical forest inventories that we used in this paper are summarized in Table 2.

**Table 2:**
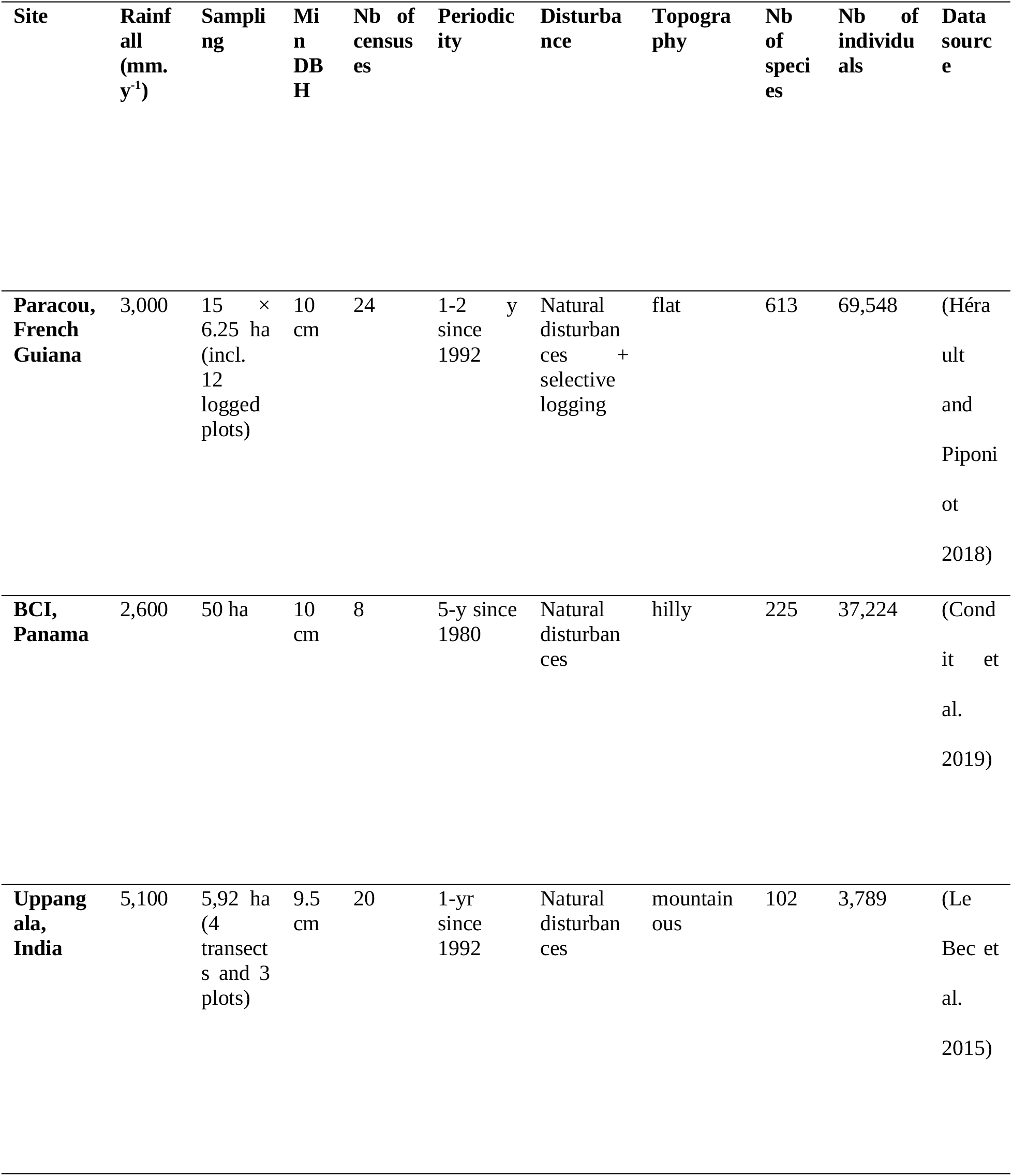
Features of the three tropical forest data sets used as empirical case studies.

For all three datasets, annualized growth between two censuses was computed as the difference of DBH (≥ 10 cm) between two consecutive censuses, divided by the time period between those two censuses. Growth estimates < -2 or > 100 mm.y^-1^ as well as individuals from incompletely identified species and individuals and species with a single observation were discarded prior to analysis. Mean annual growth for each individual tree was then computed as the difference of DBH between the first and the last time a tree was measured, divided by the time period between those two measurements.

### High observed intraspecific variability in tree growth in tropical forests

To quantify the relative importance of intra- vs. inter-specific variability in each site, we used a hierarchical growth model (Eq. IV), including an intercept *β*_*0*_, a diameter (*D*) fixed effect *β*_*1*_, a species random effect *b*_*0j*_ (with *b*_*0j*_ *∼ N(0, V*_*b*_)) and an individual random effect *d*_*0i*_ (with *d*_*0i*_ *∼ N(0, V*_*d*_)) on the intercept. All data were log-transformed and scaled, and a constant of 2 mm was added to all growth values to avoid undefined logarithms.

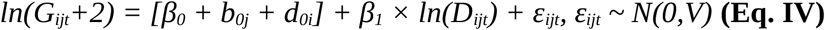

For Paracou, which has a very large dataset, we sampled 100,000 growth values randomly to perform inference. No sampling was done for Uppangala and BCI. We used priors with inverse-gamma distributions (with shape and scale parameters=10^−3^) for variance parameters, and normal distributions (with mean=0 and variance=1) for mean parameters. We estimated the inter- and intra-specific growth variability from the variance of the species (*V*_*b*_) and individual (*V*_*d*_) random effects, respectively. Model parameters were estimated the same Bayesian approach as for the analysis of the *Eucalyptus* dataset.

For the three sites, IV estimated from the growth model (*V*_*d*_, ranging from 0.41 to 0.66) was of the same order of magnitude as the interspecific variance (*V*_*b*_, ranging from 0.36 to 0.66) (Table 3). Overall, a large share of the variability in tree growth comes from individual effects in the three sites, even after accounting for the effect of diameter on tree growth, showing a high intraspecific variability in growth in these tropical forests.

**Table 3:** Mean posteriors of the tropical forest model and their estimation errors and residual variance partitioning among the different random effects.

| | Intercept ($\beta_0$) | Diameter ($\beta_1$) | Species variance ($V_b$) | Individual variance ($V_d$) | Residual variance ($V$) |
| --- | --- | --- | --- | --- | --- |
| **Paracou** |  |  |  |  |  |
| Estimate | 6.70E-02 | 2.90E-02 | 4.70E-01 | 4.60E-01 | 7.60E-01 |
| Estimation error | 2.20E-02 | 3.80E-03 | 1.70E-02 | 3.90E-03 | 2.30E-03 |
| % Variance |  |  | 27.81% | 27.22% | 44.97% |
| **Uppangala** |  |  |  |  |  |
| Estimate | 8.40E-02 | 1.90E-01 | 3.60E-01 | 6.60E-01 | 5.90E-01 |
| Estimation error | 4.40E-02 | 1.20E-02 | 4.30E-02 | 8.60E-03 | 1.90E-03 |
| % Variance |  |  | 22.36% | 40.99% | 36.65% |
| **BCI** |  |  |  |  |  |
| Estimate | 1.90E-01 | -2.20E-02 | 6.60E-01 | 4.10E-01 | 8.10E-01 |
| Estimation error | 5.00E-02 | 4.50E-03 | 3.50E-02 | 4.00E-03 | 2.00E-03 |
| % Variance |  |  | 35.11% | 21.81% | 43.09% |

### Spatial autocorrelation of individual growth within species at the local scale in tropical forests

To test whether individual growth was spatially autocorrelated, we performed in each site, spatial analyses of the mean individual growth values. We chose a conservative approach based on mean individual growth without accounting for the effect of diameter, thus without removing ontogenetic differences and considering the pattern of individual growth as it is in the field. More specifically, we performed Moran’s I one-tailed tests as implemented in the *ape* R package (Paradis and Schliep 2019), for pairs of conspecifics less than 100 m apart in the same plot (to avoid capturing the effect of treatment in Paracou and including the spaces between the plots). For the most abundant species, we sampled 3,000 individuals with a uniform probability. We considered only the species with more than five conspecific neighbors less than 100 m-apart in the same plot.

Positive spatial autocorrelation in tree growth between conspecifics was significant for 19 to 31% of the species in the three sites, representing between 45 and 79% of the total number of individuals (Table 4). Spatial autocorrelation was however much higher in logged plots as compared to unlogged in Paracou, because of a more heterogeneous light environment resulting from logging history (Appendix 3).

**Table 4:** Spatial autocorrelation of the growth of conspecific individuals in three tropical forest sites. Shown are the proportion of species, and of corresponding individuals, in percent, for which individual growth among conspecific individuals is significantly positively spatially autocorrelated. The spatial autocorrelation of individual growth was tested using Moran’s I index.

|  | Significant | Not significant |
| --- | --- | --- |
| **Paracou** |  |  |
| % Species | 31.00 | 69.00 |
| % Individuals | 78.90 | 21.10 |
| **Uppangala** |  |  |
| % Species | 18.50 | 81.50 |
| % Individuals | 45.30 | 54.70 |
| **BCI** |  |  |
| % Species | 20.10 | 79.90 |
| % Individuals | 54.70 | 45.30 |

### Higher similarity of growth within conspecific-than heterospecific individuals locally in tropical forests

To test if the performance of conspecific individuals was locally more similar than the performance of heterospecific individuals in the three sites, we also used mean individual growth, thus ignoring ontogenetic differences. We computed the mean individual growth semivariance (Baraloto and Couteron 2010) considering either conspecific or heterospecific neighbors within a 100-m radius. In the first case, semivariance was estimated as the mean of the squared difference in individual mean growth for all pairs of conspecific individuals. In the second case, semivariance was estimated as the mean of the squared difference in individual mean growth for all pairs of individuals with an individual of the focal species and one of another species. We considered only the species with more than five individuals, and with more than five heterospecific neighbors within the 100-m neighbohood distance. For each species, we then compared the semivariances between conspecific and heterospecific individuals using a Mann-Whitney test with a 0.05 alpha-risk.

The mean individual growth semivariance appeared significantly higher among heterospecifics than among conspecifics for 42 to 61% of the species in the three sites, representing 58 to 89% of the total number of individuals (Table 5). To control for a potential effect of species abundance on the semivariance estimations, we replicated the analysis by sampling a maximum of ten individuals per species. The results were qualitatively unchanged (Appendix 3).

**Table 5:**
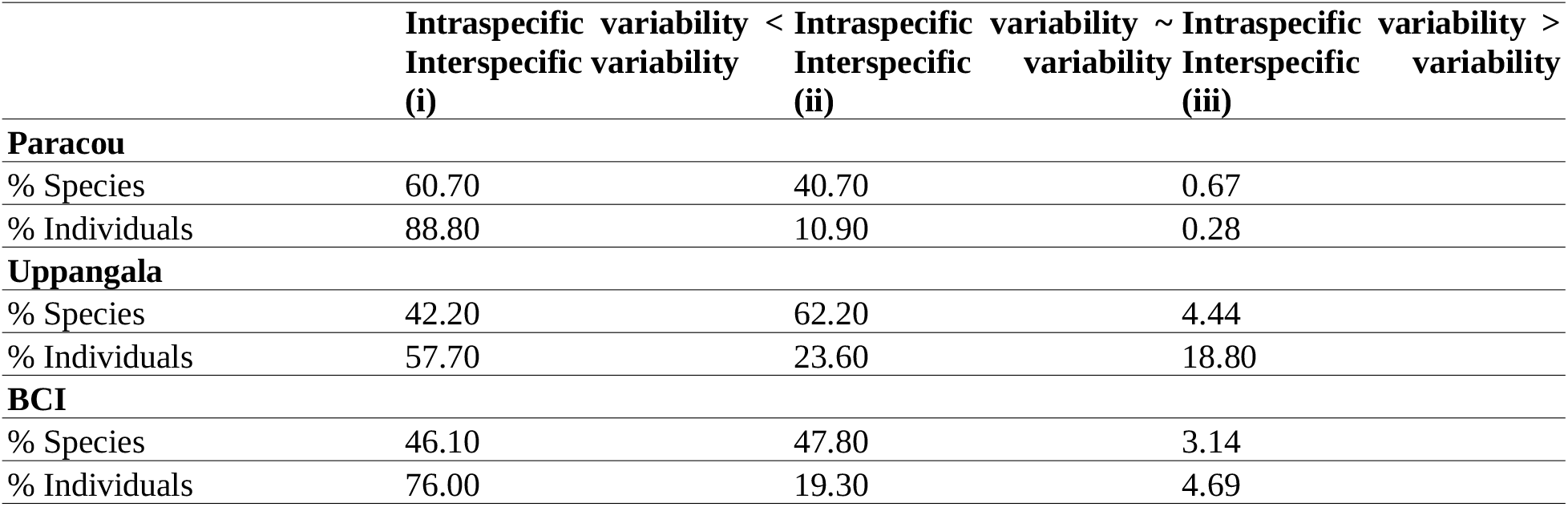
Comparison of local intra- and interspecific variability in individual growth for three tropical forest sites. The variability was estimated with the semivariance and the comparison was performed with a Mann-Whitney’s test. The semivariances were computed for all species with > 5 individuals and > 5 heterospecific neighbors within 100 m in the same plot, and considering pairs of individuals that were less than 100 m apart and in the same plot. Shown are the proportion of species, and of corresponding individuals, for which (i) intraspecific variability was significantly lower than interspecific variability, (ii) intraspecific variability was significantly higher than interspecific variability, or (iii) the difference between inter- and intraspecific variabilities was not significant.

## Discussion

### High-dimensional environmental variation leads to large observed intraspecific variability

IV can result from intrinsic differences among individuals or from extrinsic environmental variation, including biotic factors, or interactions of both (Violle et al. 2012, Moran et al. 2016, Westerband et al. 2021). While much emphasis has been placed on genetically-driven IV in studies on coexistence (Booth and Grime 2003, Ehlers et al. 2016, Barabás and D’Andrea 2016), sometimes implicitly through the use of independent random draws across individuals (Lichstein et al. 2007, Hart et al. 2016, Crawford et al. 2019), and although we acknowledge its ecological and evolutionary importance, we here argue that the importance of environmentally-driven IV in natural communities has been underestimated and has radically different consequences for species differences and community assembly. More specifically, we argue that a large part of observed IV can result from high-dimensional environmental variation in space and time.

First, using a simple simulation experiment, we illustrated how environmental variation in unobserved dimensions of the environment can produce large observed IV, although conspecific individuals are clones (Fig. 5). Similarly, the variance partitioning of individual tree growth within a common garden of *Eucalyptus* clones (le Maire et al. 2019) shows that the variance in growth between individuals is about twice as high as the variance between genotypes (Table 1). This reveals that a large part of the observed IV can emerge from environmental variables, even when the variation of the environment was sought to be minimized.

Importantly, because IV can emerge from environmental heterogeneity without underlying genetic differences, observed IV does not necessarily imply that conspecific individuals substantially differ in their response to the environment, nor that species niches overlap (Fig. 3). Instead, large observed IV can also reflect the projection of species’ high-dimensional niches within a high-dimensional environment that is variable in time and space: conspecific individuals differ with each other because they each thrive in a different micro-environment. In empirically observed data, such IV is therefore the result of projecting a high-dimensional response (*e*.*g*., physiological processes), which is controlled by multiple macro- and micro-environmental variables, down to a low-dimensional, integrative response (*e*.*g*., annual growth) that is poorly characterized because of an incomplete view of the environmental variables that contribute to it. This reassigns an important part of observed variation among individuals, often perceived as neutral or random since they are seemingly unrelated to the observed dimensions of the environment (Fig. 3, Fig. 5, Table 1), to the classical niche theory (Hutchinson 1957) and is in agreement with natural history observations of individual trait differences that are associated to species-specific ecological strategies.

The “biodiversity paradox” highlights that a large number of species can coexist while competing for a limited number of resources (Hutchinson 1961). This puzzling question has generally been tackled considering trade-offs along a limited number of niche axes, often corresponding to resources (Tilman 1982, Rees 2001). But if the number of resources may indeed be relatively limited (*e*.*g*., light, water, and nutrients for plants), the number of independent environmental factors (*e*.*g*. microclimatic variables) that drive the performance of individuals for a particular level of resources is not. Environments are known to vary along multiple dimensions at fine scales in space and time (Fig. 1), and in many cases, this variation has been shown to influence individual attributes (*e*.*g*., Fortunel et al. 2020).

Nevertheless, despite technological advances, many of these abiotic and biotic environmental factors are still poorly understood and monitored. As a result, the dimensionality of field observations is typically low compared with the high dimensionality of the environment in nature (Bramer et al. 2018, Estes et al. 2018). The variability in individual attributes due to the variation of unobserved environmental variables therefore remains mostly a black box, and is typically summarized in terms of residual variance in statistical models (Albert et al. 2012, Siefert et al. 2015) or encapsulated into so-called “individual random effects” (Clark et al. 2007). We here emphasize that even in the absence of any intrinsic differences among conspecific individuals, a large IV can emerge from the imperfect characterization of the environment (Fig. 3, Fig. 5, Table 1), which varies in a high number of dimensions (Fig. 1).

### Intraspecific variability is structured in space and time

IV has commonly been perceived and modeled through independent random draws around the species mean in community ecology studies (Lichstein et al. 2007, Courbaud et al. 2012, Hart et al. 2016, Barabás and D’Andrea 2016, Uriarte and Menge 2018). While this representation typically results from a lack of knowledge, with randomness being used as a substitute for more detailed understanding of underlying ecological processes (Clark et al. 2007), it encapsulates strong hypotheses relating to the nature of IV that are rarely discussed. In contrast, we argue here that IV is generally non-random and structured in both space and time.

At a given time, conspecific individuals that are distributed across space can strongly vary in their attributes (Violle et al. 2012, Siefert et al. 2015, Moran et al. 2016, Poorter et al. 2018). While this spatial IV has often been interpreted as random, *i*.*e*. implying that conspecific individuals can perform differently within the same environment (Fig. 3c), a large part of this variability appeared in fact structured in space and likely associated with fine-scale spatial changes in the environment (Fig. 3b, Moran et al. 2016). In our illustrative simulation experiment, the attribute of conspecific individuals varies spatially as a result of the environmental variation in space, and the spatial autocorrelation of conspecific attributes reflects the spatial autocorrelation of the environmental variables (Fig. 5, Fig. 6).

Similarly, data from three long-term forest inventory sites across the tropics revealed spatial autocorrelation in tree diameter growth of conspecific individuals (Table 3), suggesting that IV is strongly driven by the spatial variation of the environment, which is itself highly structured (Fig. 1). However, we acknowledge that genetically-driven IV can also be spatially structured, for instance via dispersal patterns or natural selection (Moran et al. 2016). We hypothesize that in that case, attributes would likely be randomly structured in space (Getzin et al. 2014) or correlated at the spatial scale of seed dispersal, typically several tens of meters in tropical forests (Clark et al. 2004, Seidler and Plotkin 2006, Muller-Landau et al. 2008), while environmental variables are typically highly spatially correlated at fine scales (*e*.*g*. meter scale, Baraloto and Couteron 2010). We also acknowledge that natural selection can happen at fine scales (Marrot et al. 2021), and could thus produce spatially structured IV due to local genetic adaptation. Nevertheless, data documenting genetic variation within species can still reveal higher similarity between conspecifics than heterospecifics locally as well as non-overlapping species niches (Schmitt et al. 2021). Importantly, any local genetic adaptation does not preclude that multidimensional environmental variations generate large observed IV that is structured in space and time and whose consequences cannot be well represented and understood using a random variation around a species mean.

In communities of sessile organisms such as trees, IV has been commonly structured in space using individual random effects, which vary among conspecific individuals but stay constant through the lifetime of individuals (Clark et al. 2007, Vieilledent et al. 2010). We here argue that while this approach can *reveal* the spatial structure of IV through inference, the use of the resulting estimated standard deviation term to introduce individual variation in simulations of community dynamics is not sufficient to *produce* a spatially structured IV, as we showed is observed in natural communities.

Similarly, individual attributes can change over time. Because individuals within a species can be measured at different points in time, as it is often the case when assembling functional trait databases for example (Zanne et al. 2009, Albert et al. 2011, Kattge et al. 2020) this can lead to an observed unstructured IV when characterized by a variance around a species mean (Fig. 3c). But a large part of this observed IV is actually structured in time and associated with temporal changes in the environment. For instance, the temporal storage effect (Chesson and Warner 1981), a well-known coexistence mechanism, structures species performance because species are able to “store” growth during favorable timespans to overcome lean times; mast-seeding or masting, which describes periodic and synchronized massive seed production of conspecific individuals, would also result in a temporally structured IV (Koenig and Knops 2005). Temporal variation in individual response within a species can typically be structured with temporal random effects (Clark et al. 2007). Temporal random effects have been used to estimate the inter-annual variability in tree growth (Metcalf et al. 2009, Fortunel et al. 2018) and fecundity (Clark et al. 2007) for example. In all those examples, temporal environmental variation affects conspecific attributes in the same way (Clark 2010).

We therefore call for a reconsideration of the nature and the way of integrating IV into models of community dynamics. When IV is modeled randomly with a variance around a species mean, it implies that conspecific individuals can perform differently in the exact same environment, thus implying intrinsic differences between conspecific individuals. This type of unstructured IV can result in an overestimated increase in species niche overlap, which blurs species differences (Fig. 3a and 3c, Stump et al. 2021). While trait heritability has rarely been considered in studies on the role of IV on coexistence (but see Barabás and D’Andrea 2016), in some studies, the random variation in attributes across conspecific individuals is considered as environmental, because it is not heritable in the model (*e*.*g*. Lichstein et al. 2007, Moran et al. 2016). However, environmentally-driven IV should be structured in space and time, as the environment is (Fig. 1, Fig. 6). In addition, when IV is randomly distributed among conspecific individuals, similarity among conspecific individuals is systematically underestimated, which is not the case when IV is structured in space and time (Purves and Vanderwel 2014, Banitz 2019), as discussed hereafter.

### Conspecific individuals respond more similarly than heterospecific individuals locally

Species differ in multiple attributes, responding to a high number of environmental variables (Fig. 2), but often in ways that cannot be readily observed. If observed IV results mainly from high-dimensional environmental variation in space and time rather than from intrinsic differences between conspecific individuals, then for a given environment, conspecific individuals should respond more similarly than heterospecific individuals. This is the case in our illustrative simulation experiment, where the fact that conspecific individuals have exactly the same set of parameters and respond identically to spatial and temporal changes in the environment results in higher inter-than intraspecific variance in the response locally (Fig. 5, Fig. 6).

Corroborating this point of view, pairs of conspecific individuals in 11 North-American temperate forest stands showed higher correlation in their temporal variation of growth rate or fecundity than pairs of heterospecific individuals on average (Clark 2010). This indicates that conspecific individuals responded more similarly to environmental variation in time than individuals of different species. Importantly, these results were obtained in a system with high observed IV (leading to an apparent species niche overlap), where species responded in the same direction to environmental changes (*e*.*g*. increased tree growth in climatically favorable years). Hence, considering the temporal structure of IV revealed species differences that were not apparent otherwise, since they led to spreading along a high number of dimensions that varied at fine scales (Clark 2010). However, as well highlighted by Stump et al. 2021, these results were often misinterpreted as an evidence that IV fostered coexistence. As another piece of evidence presented here, pairs of spatially proximal conspecific individuals tended to present more similar temporal means in absolute tree growth than pairs of close heterospecific individuals across three large contrasted tropical forest sites (Table 5). This provides new empirical evidence that, although estimated IV can be substantial, conspecific individuals respond more similarly than heterospecific individuals to environmental variation in space.

A stronger similarity in the response to environment between conspecific than heterospecific individuals locally leads to a stronger concentration of competition within species, which, ultimately, can result in intraspecific competition being greater than interspecific competition, a common driver of stable species coexistence (Lotka 1925, Volterra 1926, Chesson 2000). Because species differ in their response to the environment, environmental variation in space and time leads to local or punctual inversions of species hierarchy in performance (Fig. 3d). As possibilities of hierarchy inversions between species increase rapidly with increasing dimensionality of the environment (Fig. 3b), the high-dimensionality of the environment offers room for the stable coexistence of numerous species (Falster et al. 2017, Rüger et al. 2018). In the end, we therefore argue that a substantial part of IV is not a mechanism for coexistence in itself but can rather be the signature of species differences and environmental variation that allow coexistence: the high-dimensional species differences, which make them respond differently in a high-dimensional environment varying in space and time, can only be observed at the individual scale. In the absence of precise information on the many dimensions across which species differ and environment varies, large observed IV is the evidence of the niche mechanisms enabling species coexistence.

### Recommendations and concluding remarks

Most of the theoretical studies that have explored the role of IV in species coexistence so far did so by adding variances around species-specific means, thus considering IV as stochastic, which implies that conspecific individuals perform differently in the same environment. Here, we provide insights suggesting that large observed IV can emerge from environmental heterogeneity and is structured in space and time. We stress that this interpretation has strong consequences on the understanding of the effects of IV on species coexistence: (i) observed IV does not necessarily imply that conspecific individuals are strongly intrinsically different nor that species niches overlap, and (ii) the spatial and temporal structure of observed IV reveals stronger concentration of competition within species locally in space and time, which is a frequent necessary condition for stable species coexistence. We thus call for a reconsideration of the nature of IV and of the way it is integrated in models, by thoroughly distinguishing its sources (intrinsic vs. extrinsic, and their interactions). We acknowledge the existence of genetically-driven IV, potentially due to local adaptation to the microenvironment, and its eco-evolutionary importance, but suggest that multidimensional environmental variation generates a large observed IV that is structured in space and time. We underline that environmentally-driven structured IV has been largely overlooked in previous community ecology studies and has consequences on community dynamics which cannot be represented and understood using a random variation around a species mean. To this end, we recommend that empirical studies explore further the spatio-temporal structure of IV and how it relates to environmental variation along multiple dimensions, and, when possible, assess the relative importance of genetically and environmentally driven IV, for instance by means of common garden experiments. Models of community dynamics should then endeavor to structure IV in space and time so that it reflects the high-dimensional variation in both the environment and species attributes, and not only some intrinsic differences between conspecific individuals (Purves and Vanderwel 2014, Moran et al. 2016, Banitz 2019). In both empirical studies and models, this implies that the species attributes are measured at the individual level, localized in space, and repeatedly observed in time. Simultaneously, the monitoring of multiple environmental variables at fine scales in space and time is required in order to better capture their effect on individual attributes (such as physiological or mechanistic traits, Shipley et al. 2016, Brodribb 2017), hence reducing the part of unexplained IV, and ultimately to better characterize the high-dimensionality of species niches. Altogether, these recommendations will enable to better account for species differences that are expressed at the individual level and evidence their impacts on the community dynamics in natura and in silico.

## Supporting information

Complete Supplementary Information

## Acknowledgments

This paper is a joint effort of the working group “INTRACO” supported by CESAB, the synthesis center of the French Foundation for Research on Biodiversity (FRB) and sDiv, the synthesis center of iDiv (DFG FZT 118, 202548816). CGT’s work is supported by a PhD grant provided by the Laboratoire d’Excellence CEBA (Center for the study of Biodiversity in Amazonia; http://www.labex-ceba.fr/en/). CEBA is funded by an “Investissement d’Avenir’’ grant of the French National Research Agency (CEBA: ANR-10-LABX-25-01). Data supporting our point of view have been collected by the field teams of Paracou (CIRAD), the EUCFLUX project, Uppangala (Karnataka Forest Department and French Institute of Pondicherry), and Barro Colorado Island (Smithsonian Tropical Research Institute). Authors thank Kasey Barton and Sean McMahon for helpful discussions and comments on previous versions of the manuscript.

## Supplementary information and data access

The appendices and all the code used for this study are available in a GitHub repository (https://github.com/camillegirardtercieux/coexIV) and have been permanently archived on Zenodo (https://doi.org/10.5281/zenodo.5504013).

Appendix 1: Simulation experiment with two species.

Appendix 2: Analysis of an *Eucalyptus* clonal plantation dataset.

Appendix 3: Analysis of tropical forest inventory data.

No new data were used in this study. For access to forest plot inventory data and *Eucalyptus* plantation data used in this study, refer to the data used in Le Bec et al. 2015, Hérault and Piponiot 2018, Condit et al. 2019 and le Maire et al. 2019. However, the analyses and reflections presented here are original.

## Statements of author roles

*CGT, IM and GV conceived the initial ideas and coordinated the INTRACO working group. All authors contributed to the study design and ideas within the INTRACO working group. CGT led the analyses. CGT, IM and GV wrote the first draft of the manuscript, and all authors contributed substantially to revisions*.

