## Supplementary material for "Rethinking the nature of intraspecific variability and its consequences on species coexistence": Complete Supplementary Information

**References** **34**

### 1 Appendix 1: simulation experiment with two species

#### 1.1 Perfect knowledge model: generating response values.

We fixed the species parameters of the "Perfect knowledge model" as follows (see main text, Eq. 1), using 10 environmental variables ($N$=10):

Parameters of species 1: $\beta_0 = 0.25$, $\beta_1 = 0.15$, $\beta_2$ to $\beta_6$ were chosen randomly between -0.05 and 0.05 and $\beta_7$ to $\beta_{10}$ were chosen randomly between 0 and 0.05.

Parameters of species 2: $\beta_0 = 0.2$, $\beta_1 = 0.1$, $\beta_2$ to $\beta_{10}$ were chosen the same way as for species 1.

The difference between the species was imposed by those parameters. Here, species 1 was more competitive on average thanks to its higher intercept ($\beta_0$) and response to the first environmental variable ($\beta_1$) (see main text, Fig. 3). The first environmental variable ($X_1$) had a higher weight in the computation of the response variable, as would be the most limiting factor for the response variable.

To account for intrinsic variability in our generated data, we added variability in species parameters by sampling each individual parameter in a normal distribution centered on the species mean parameter and with a standard deviation of a quarter of the species mean parameter.

To represent the spatialized environment, we built a 2D matrix of dimension 500 × 500 for each environmental variable at a time $t_0$, by randomly generating them with spatial autocorrelation. Each variable was independently simulated by using the gstat R package (Pebesma 2004, Gräler et al. 2016), enabling to create autocorrelated random fields. A spherical semivariogram model was used for each of the ten environmental variables, with a mean of 0 for each explanatory variable (beta = 0), a sill of 1 (psill = 1) for each and a range of 50 (range = 50).

We then considered that $Y$ had been measured at two times, $t_0$ and $t_1$, for each individual. A pair of coordinates within this spatialized environment was randomly assigned to each of the 500 individuals. We considered that some of the environmental variables (light, temperature, humidity, nutrient availability for instance) had changed between $t_0$ and $t_1$ and others had not (slope, altitude for instance). For the first environmental variable which had a stronger impact on $Y$ values and another randomly drawn environmental variable, we computed values at $t_1$ as $t_0 + \epsilon$, $\epsilon \sim \mathcal{N}(0, 0.1)$ (they randomly increased or decreased). For two other environmental variables randomly drawn, $X_{t_1} = X_{t_0} + |\epsilon|, \epsilon \sim \mathcal{N}(0, 0.1)$ (they increase) and for two other environmental variables, $X_{t_1} = X_{t_0} - |\epsilon|, \epsilon \sim \mathcal{N}(0, 0.1)$ (they decreased).

This led to two repeated measurements of $Y$ for each individual of each species, i.e. 2000 values of $Y_{i,j,t}$, with $i = [1; 2]$ ; $j = [1, 50]$ ; $t = [t_0; t_1]$.

Figure S1.1 shows the distribution of the simulated $Y$ data and figure S1.2 shows the relationship between $X_1$ and $Y$.

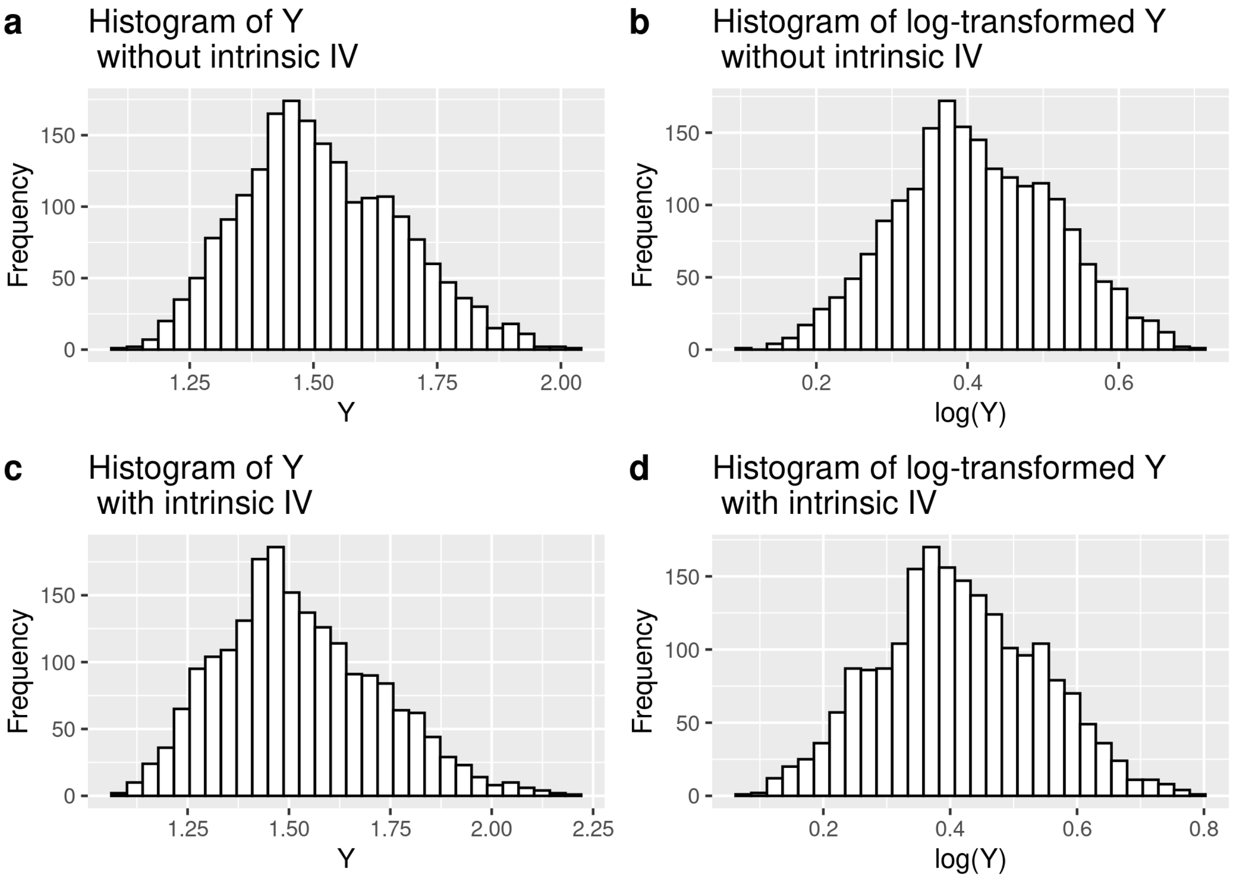

Figure S1.1: Histograms of the simulated $Y$ values (a) without intrinsic IV (b) without intrinsic IV and log-transformed (c) with intrinsic IV (d) with intrinsic IV and log-transformed.

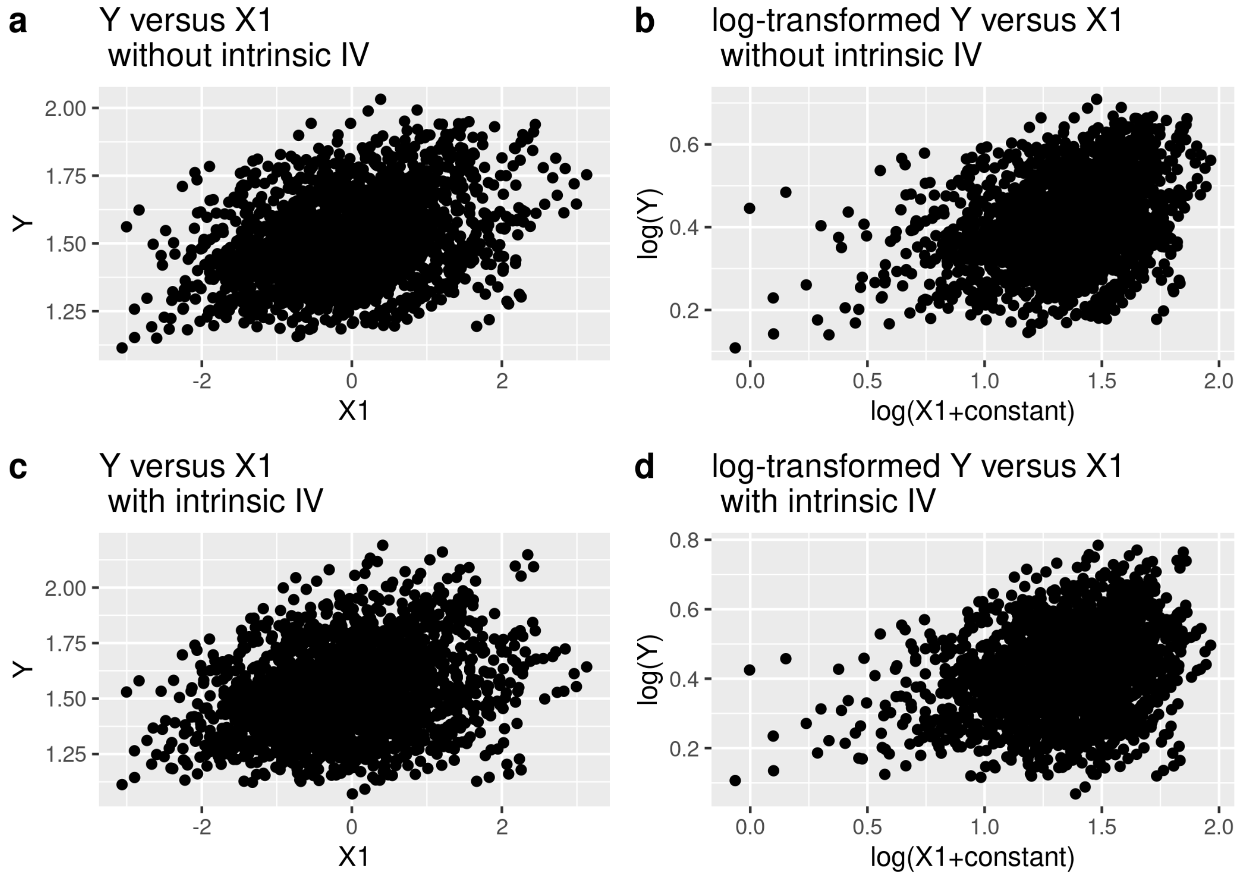

Figure S1.2: Raw and log-transformed $Y$ versus $X_1$ plots with and without intrinsic IV.

#### 1.2 Imperfect knowledge model: interpreting the response values with only one explanatory variable.

We ran the model (see main text, Eq. 2) twice, with the datasets generated with and without intrinsic IV. We visualized the convergence and the results of the models thanks to trace and density plots.

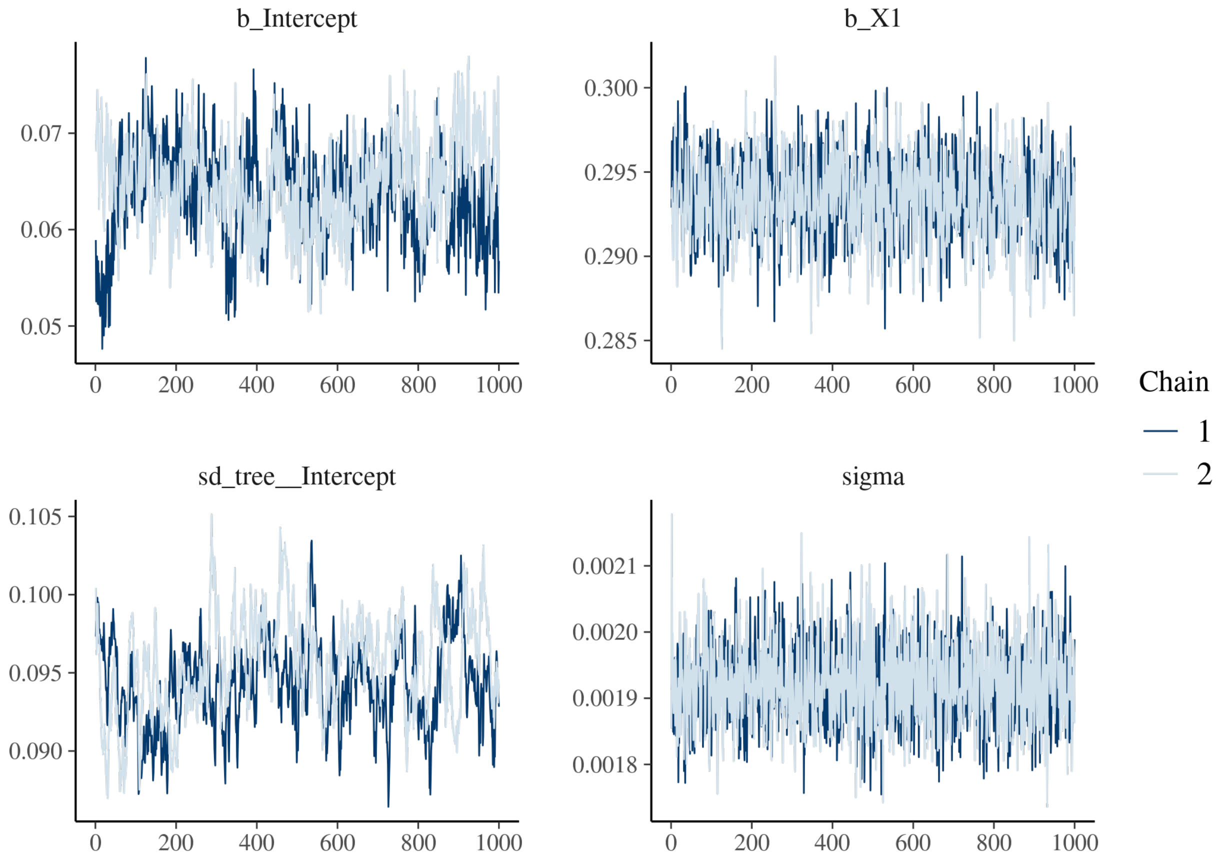

Figure S1.3: Trace of the posterior of the model for Species 1 without intrinsic IV

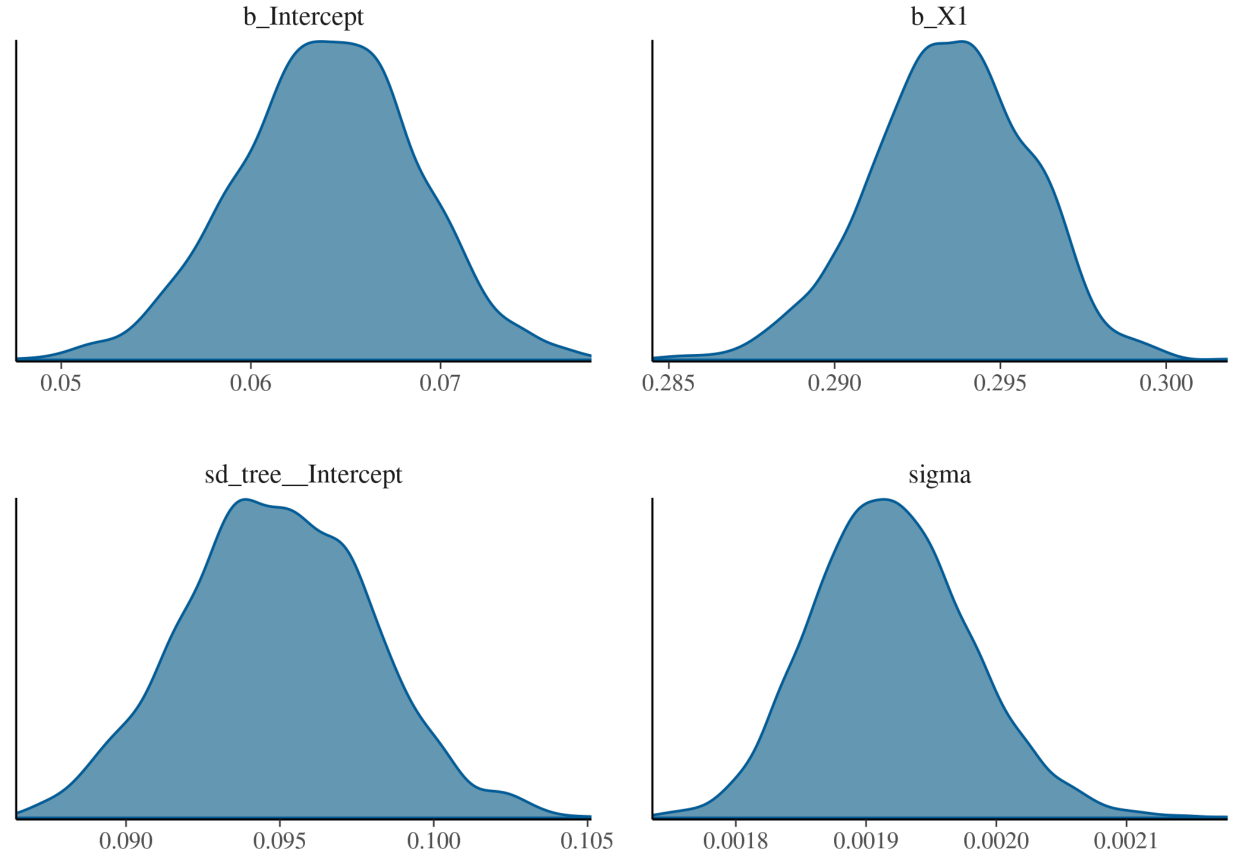

Figure S1.4: Density of the posterior of the model for Species 1 without intrinsic IV

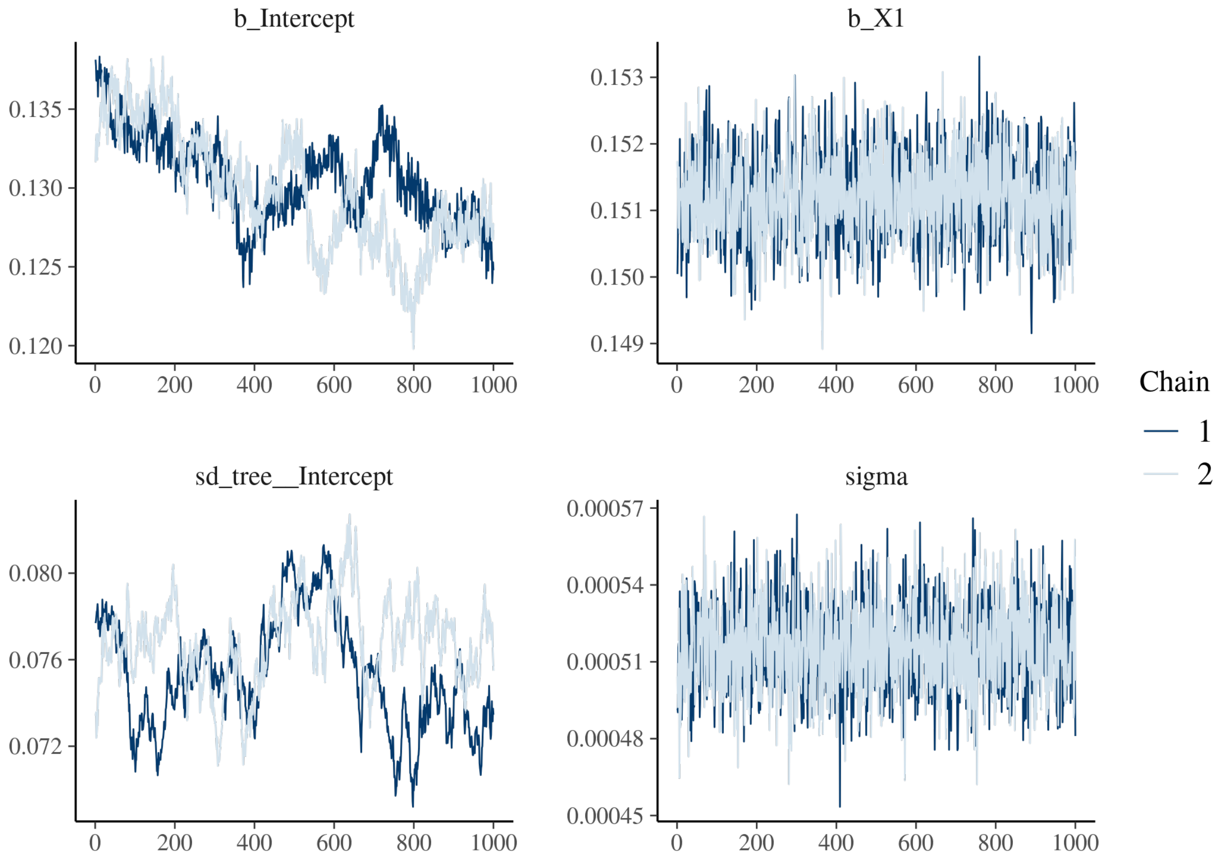

Figure S1.5: Trace of the posterior of the model for Species 2 without intrinsic IV

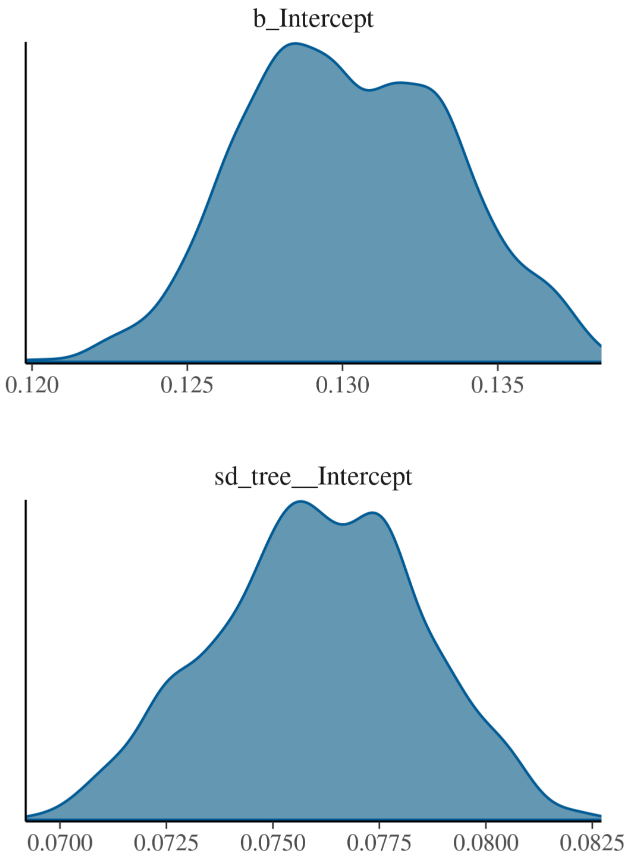

Figure S1.6: Density of the posterior of the model for Species 2 without intrinsic IV

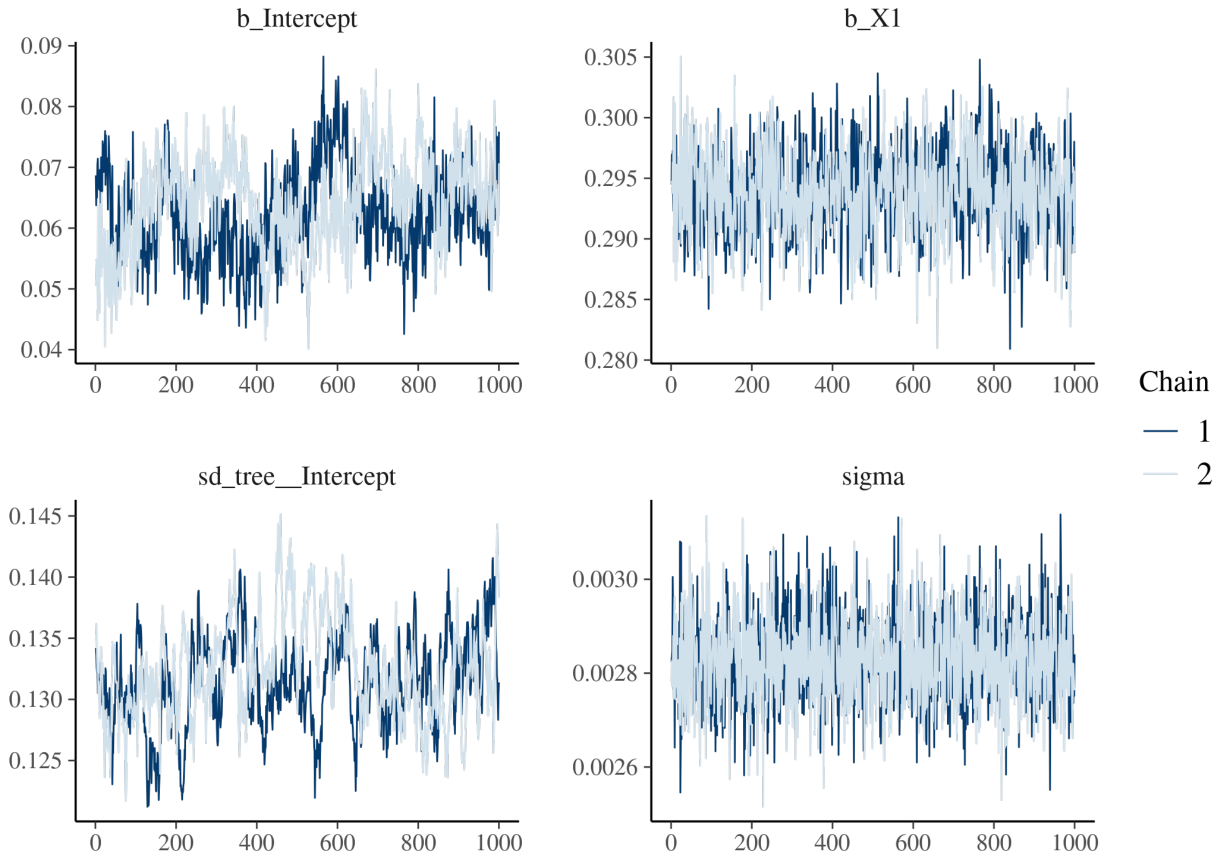

Figure S1.7: Trace of the posterior of the model for Species 1 with intrinsic IV

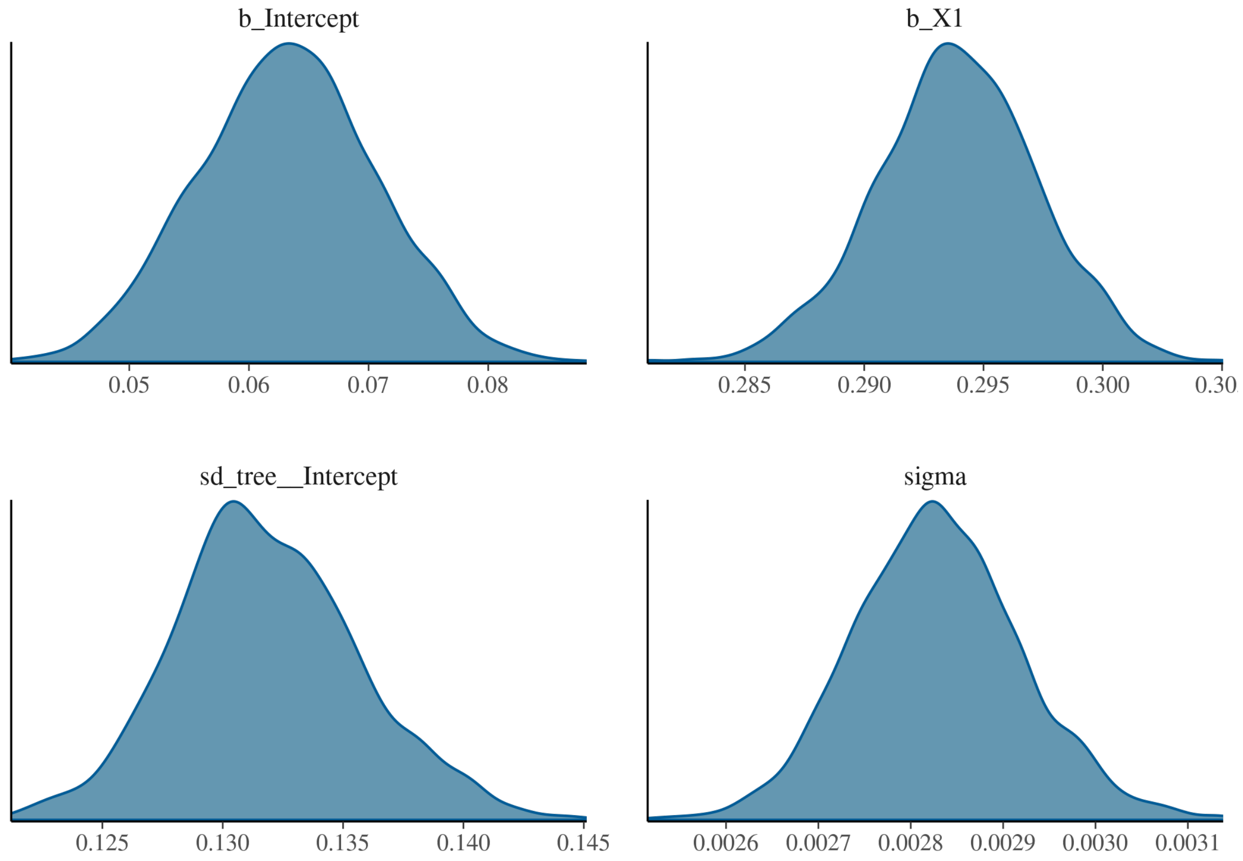

Figure S1.8: Density of the posterior of the model for Species 1 with intrinsic IV

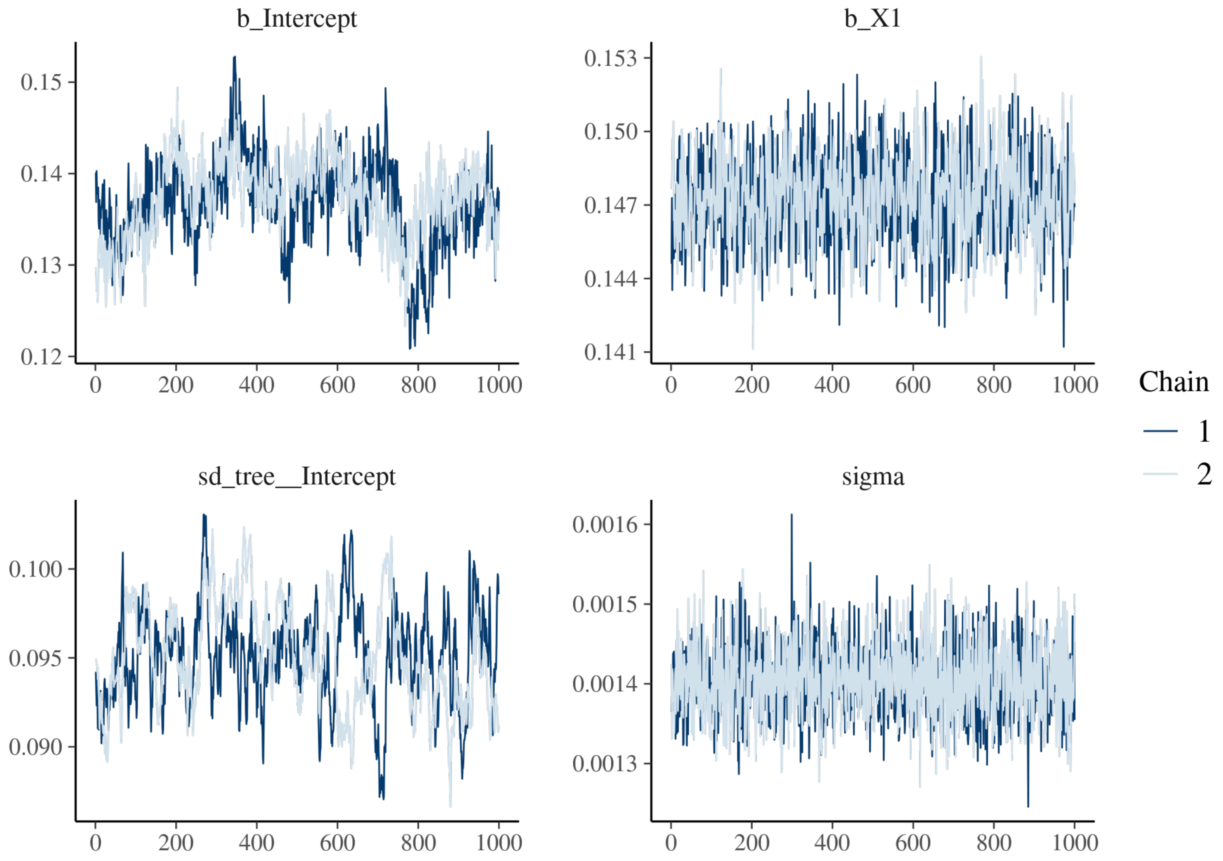

Figure S1.9: Trace of the posteriors of the model for Species 2 with intrinsic IV

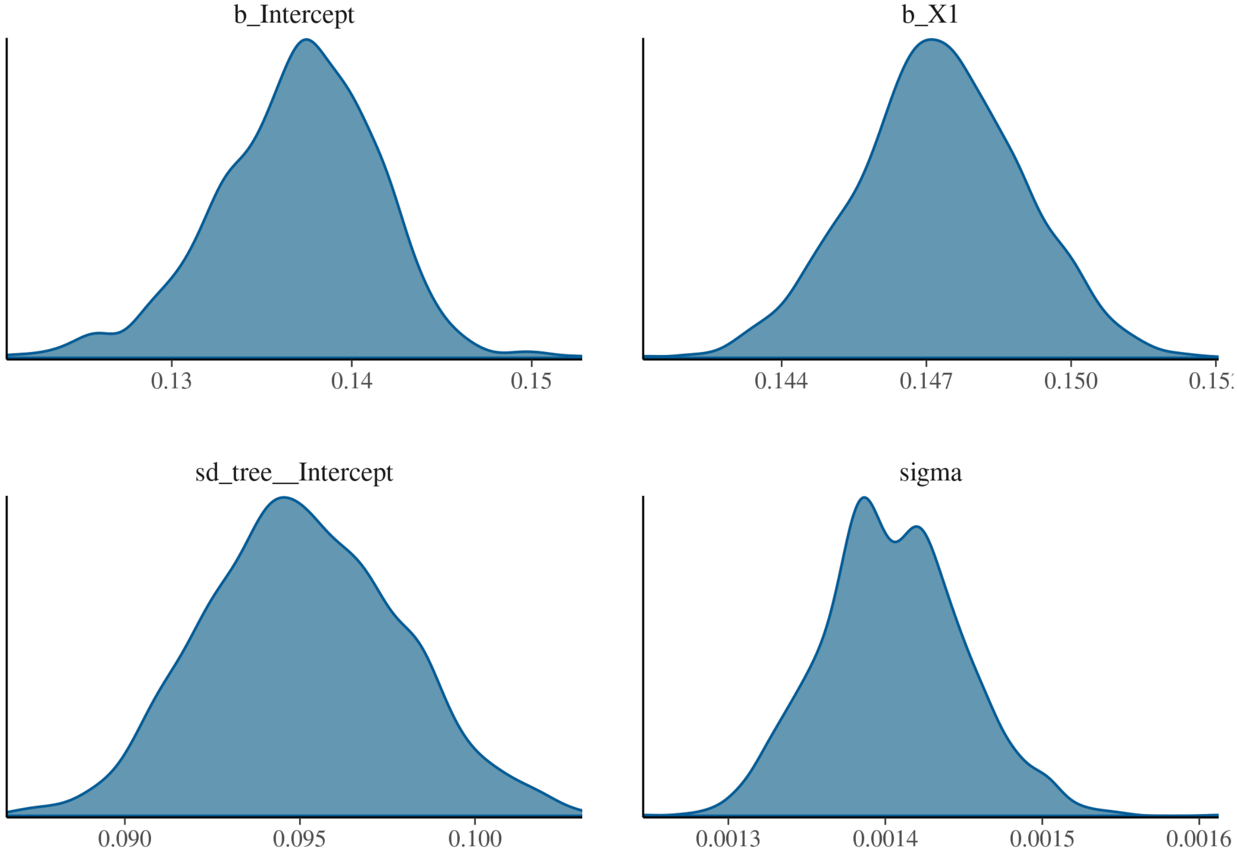

Figure S1.10: Density of the posteriors of the model for Species 2 with intrinsic IV

Table S1.1: Mean posteriors and their estimation error

| | $\beta_0$ | $\beta_1$ | $V_b$ | $V$ |
|---|---|---|---|---|
| **Species 1 (no random IV)** | | | | |
| Estimate | 6.4e-02 | 2.9e-01 | 9.5e-02 | 1.9e-03 |
| Estimation error | 4.7e-03 | 2.4e-03 | 3e-03 | 6.1e-05 |
| **Species 2 (no random IV)** | | | | |
| Estimate | 1.3e-01 | 1.5e-01 | 7.6e-02 | 5.2e-04 |
| Estimation error | 3.3e-03 | 6.3e-04 | 2.4e-03 | 1.7e-05 |
| **Species 1 (with random IV)** | | | | |
| Estimate | 6.3e-02 | 2.9e-01 | 1.3e-01 | 2.8e-03 |
| Estimation error | 7.4e-03 | 3.4e-03 | 3.9e-03 | 9.2e-05 |
| **Species 2 (with random IV)** | | | | |
| Estimate | 1.4e-01 | 1.5e-01 | 9.5e-02 | 1.4e-03 |
| Estimation error | 4.5e-03 | 1.7e-03 | 2.8e-03 | 4.5e-05 |

We inferred a high IV even in the absence of intrinsic IV (Table S1.1).

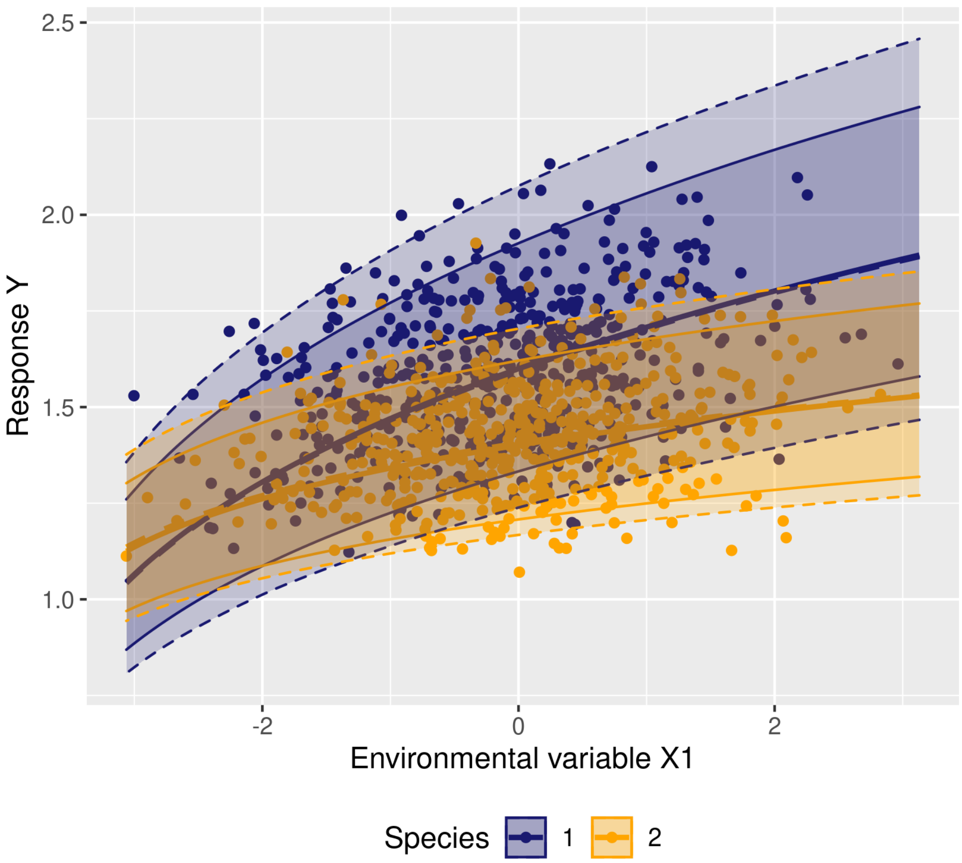

Figure S1.11: Plot of the real values - points - and estimated mean - bold lines - and 95% confidence interval - thin lines - of $Y$ versus $X_1$. The dashed lines correspond to the 95% interval due to intrinsic IV.

In Figure S1.11 the bold and solid (dashed) lines represent the mean rate of the response variable (e.g. growth) of Species 1 (blue) and Species 2 (orange) as computed with the parameters retrieved from the model without (with) intrinsic IV respectively. The plain lines represent the 95% interval of the posteriors from the model without intrinsic IV and the dashed lines show the 95% confidence interval of the posteriors from the model with intrinsic IV. Figure S1.11 shows that intrinsic IV simply increases the overlap between the response of Species 1 and Species 2.

#### 1.3 Spatial autocorrelation of individual response

To test for spatial autocorrelation of individual response, we computed Moran's I test using the Moran.I function of the ape R package (Paradis and Schliep 2019).

Table S1.2: Results of the Morans's I test for both species separatly and together.

| | Moran's I index | P-value |
|---|---|---|
| Species 1 | 6.7e-02 | 0e+00 |
| Species 2 | 6.9e-02 | 0e+00 |
| All individuals | 4.3e-02 | 0e+00 |

##### 1.4 Similarity between conspecific individuals compared to heterospecific individuals and consequences for species coexistence.

We used semivariograms to visualise the spatial autocorrelation of the response variable and to test whether the individual response was more similar within conspecifics than within heterospecifics. The semivariograms were computed and modelled with the variogram and the fit.variogram functions of the gstat R package (Gräler et al. (2016)) respectively. The variogram models were spherical.

#### 2 Appendix 2: analysis of an Eucalyptus clonal plantation dataset

##### 2.1 Data and preliminary analysis

The experimental design aimed at minimizing environmental variations and selected productive genotypes able to accommodate several environmental conditions (Maire et al. 2019).

Mean annual growth of each tree in mm/y was computed as the DBH (Diameter at Breast Height) difference between two consecutive censuses divided by the time between the two censuses. In case of mortality of the tree between two censuses, the data was discarded. We computed the neperian logarithm of diameter and growth (with a constant for growth in order to avoid undefined values).

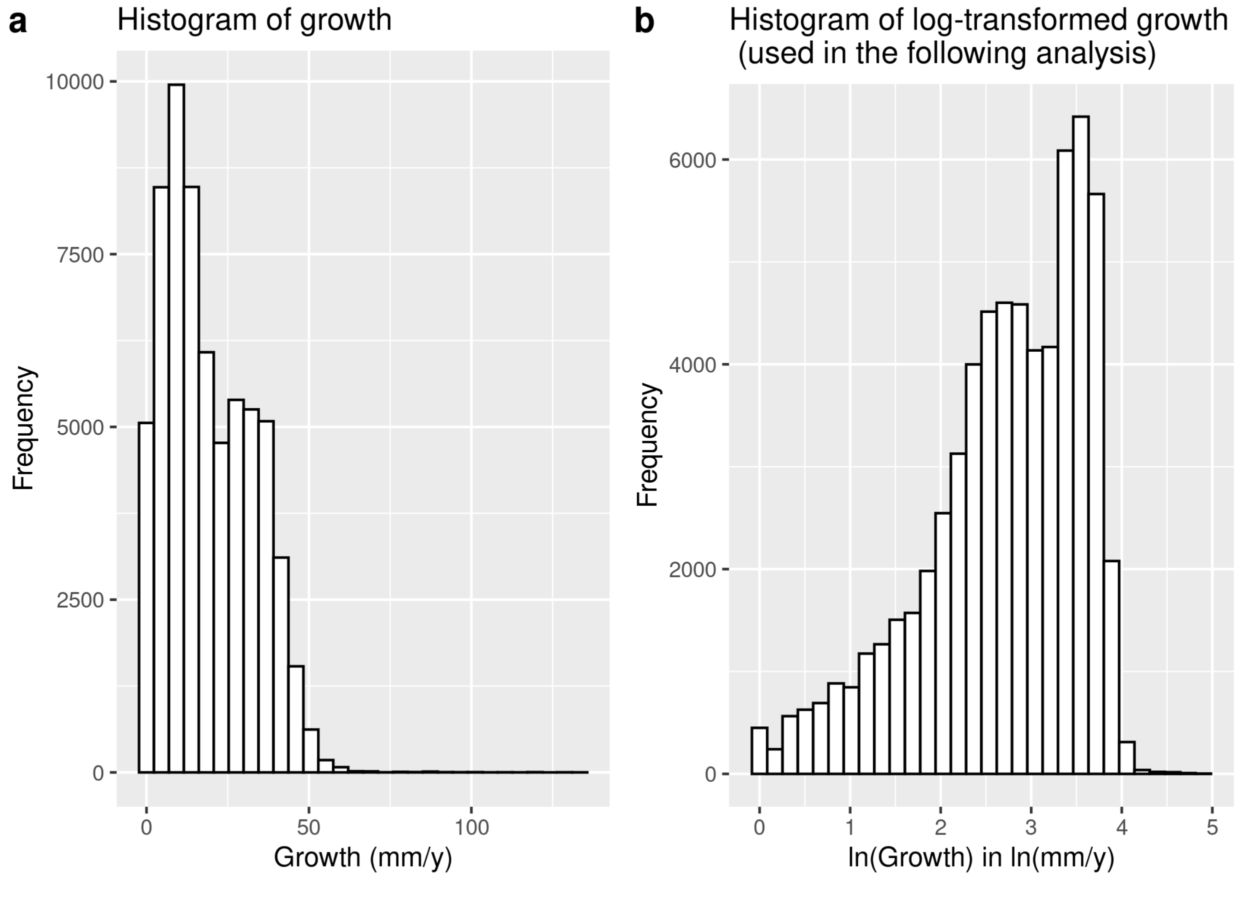

Figure S2.1: Growth (after removing data after mortality) before (a) and after (b) log-transformation.

Figure S2.1 shows the distribution of growth data before and after log-transformation.

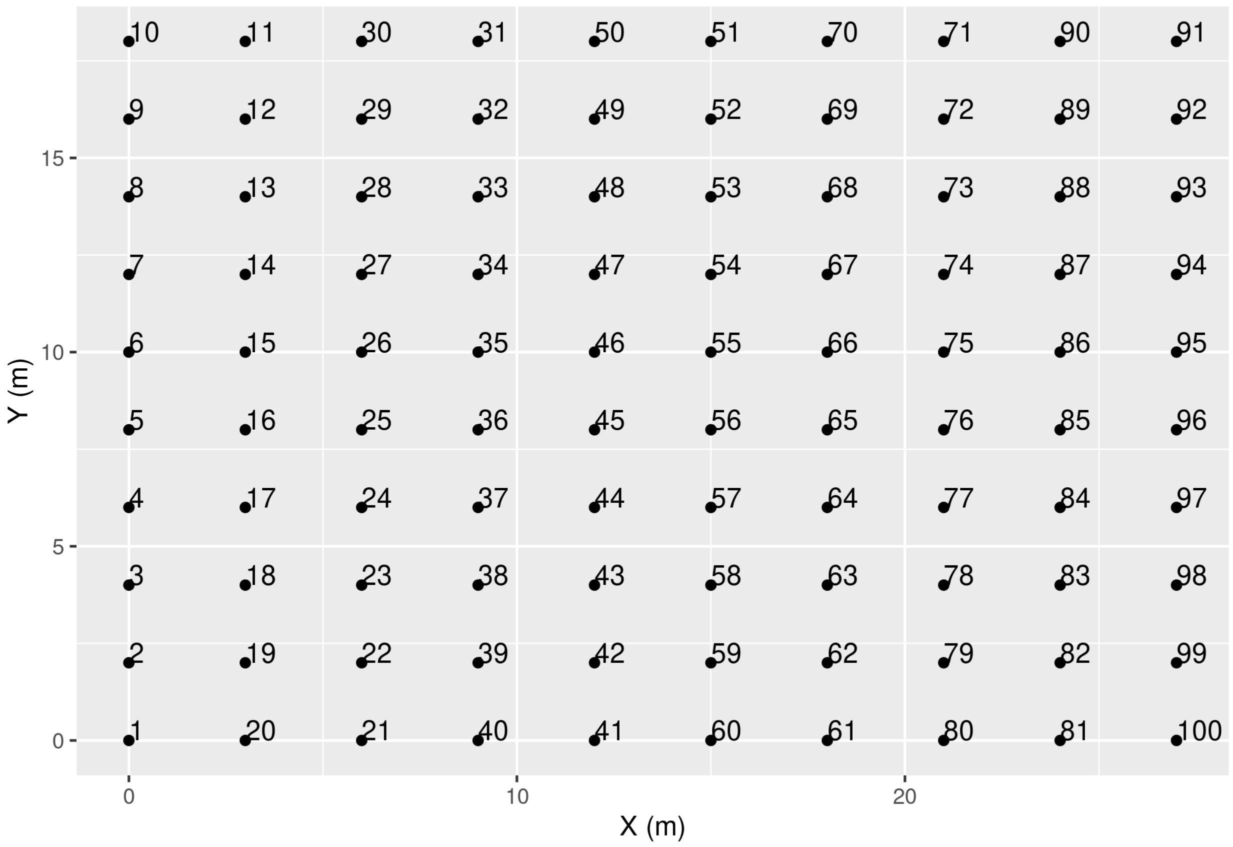

Figure S2.2: Design of a plot. Each point is a tree and the associated number is the tag of the tree.

Figure S2.2 shows the disposition of the trees in a single plot. There are 14 genotypes times 10 repetitions, so 140 plots with this same design.

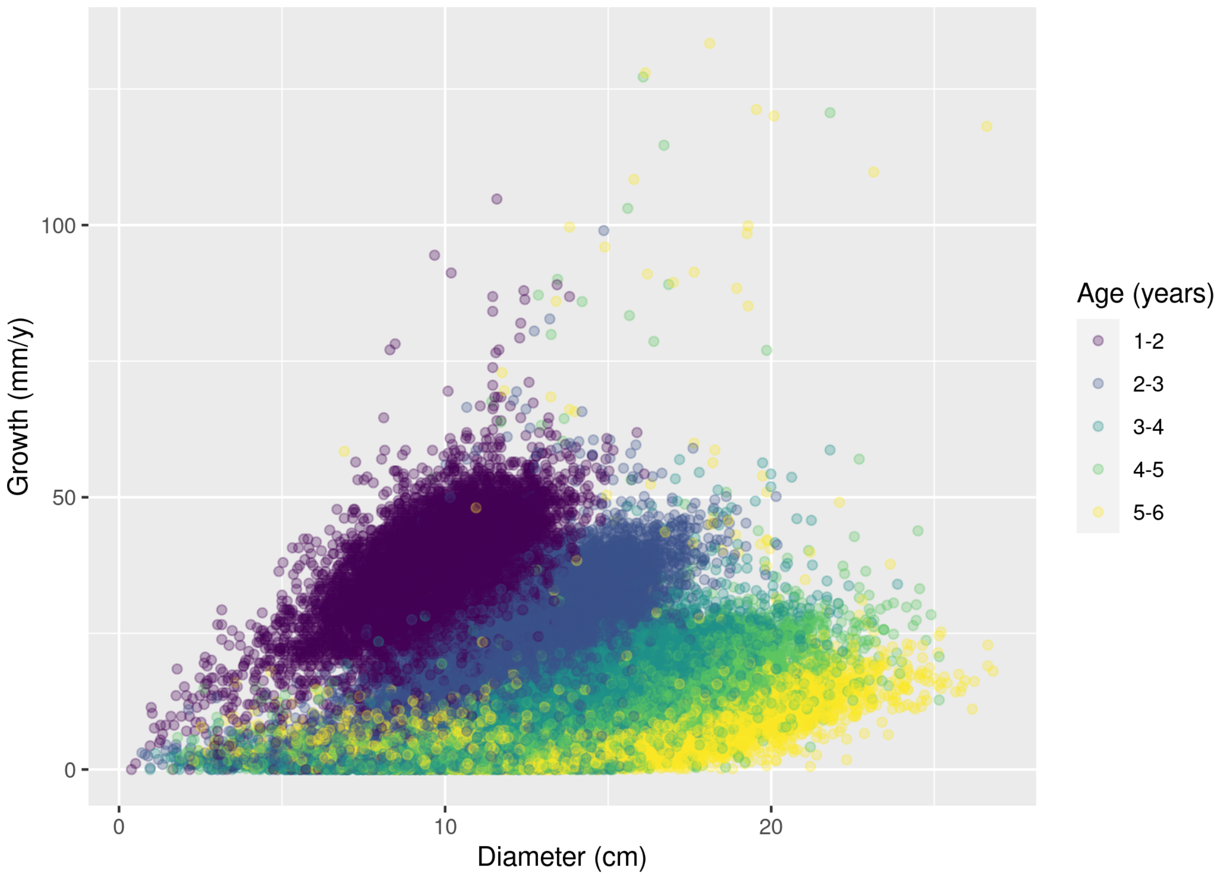

Figure S2.3: Plot of the growth versus the diameter. Each colour represents a tree age.

Figure S2.3 shows the age of the trees has a big influence on the values of growth but also on the relationship between growth and diameter: the slope is smaller with time, indicating that for the same diameter, growth is slower through time. This is likely an effect of competition for light and possibly underground resources, since as the trees grow their capacity to capture resources increases. Therefore, we computed a competition index $C_{i,t}$ to integrate this effect in the growth model. The competition index was computed for each tree which is not on the edge of a plot. It was the sum of the basal areas (BA) of the 8 direct neighbours (there was no need to divide by the area of the rectangle that comprises all the direct neighbours, since this latter is a constant by construction of the experimental design). It was then log-transformed. Dead neighbours were considered as having a null BA.

$C_{i,t} = \sum BA_{neighbours(i,t)}$

#### 2.2 Estimation of intraspecific variability

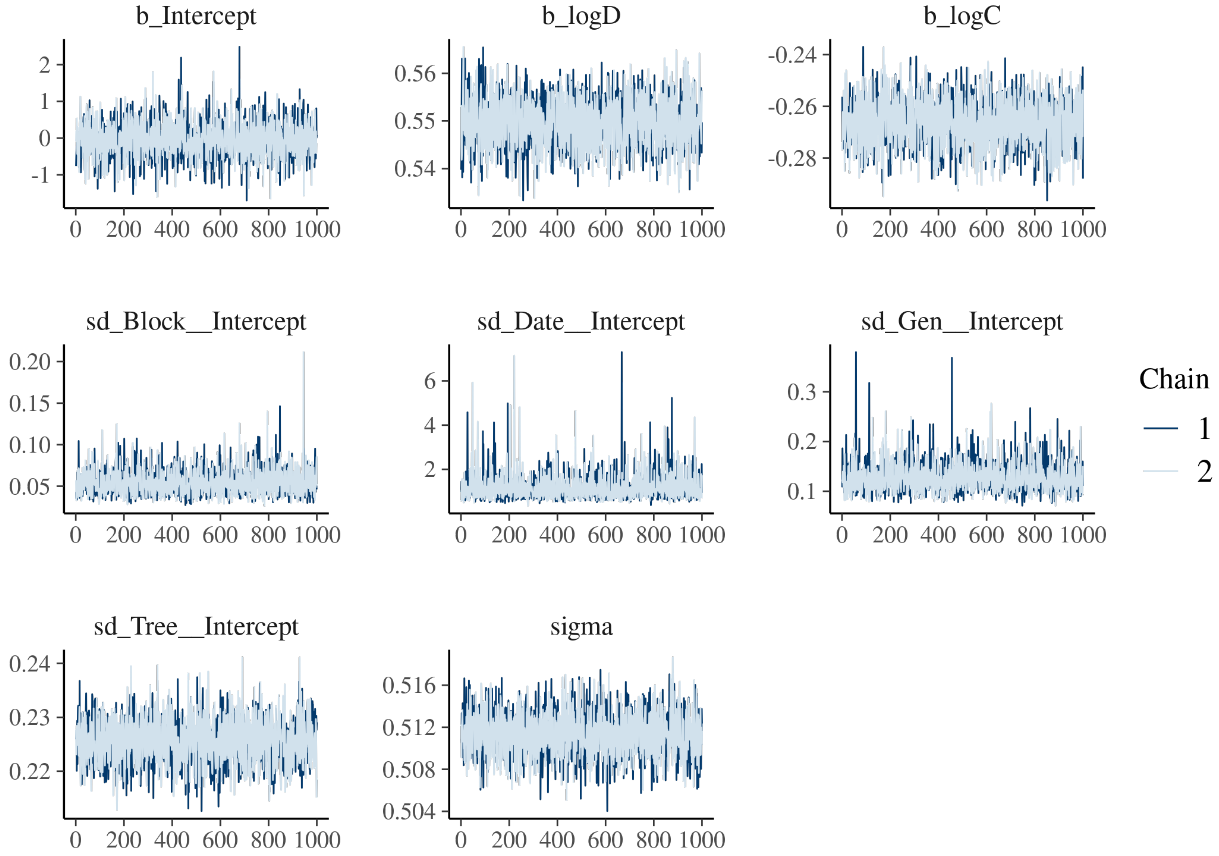

Figure S2.4: Trace of the posteriors of the inferred parameters

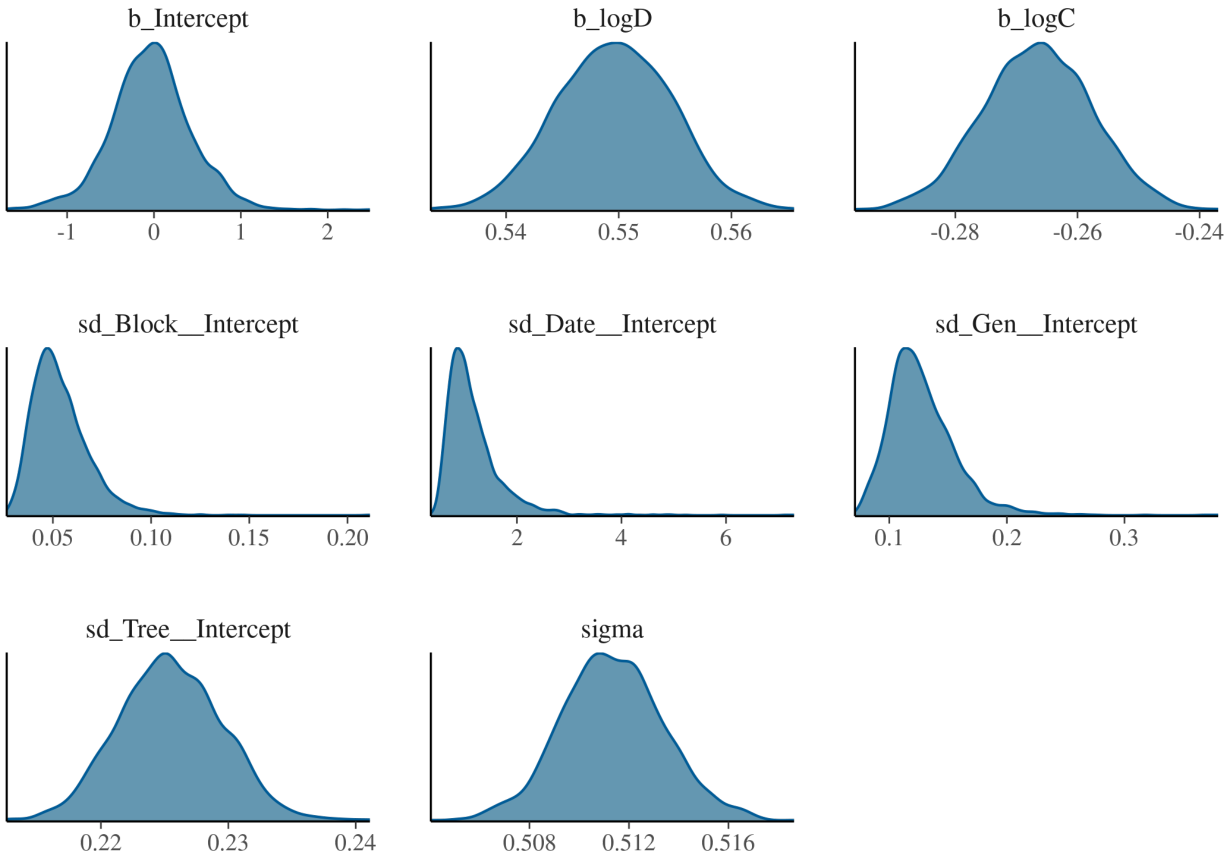

Figure S2.5: Density of the posteriors of the inferred parameters

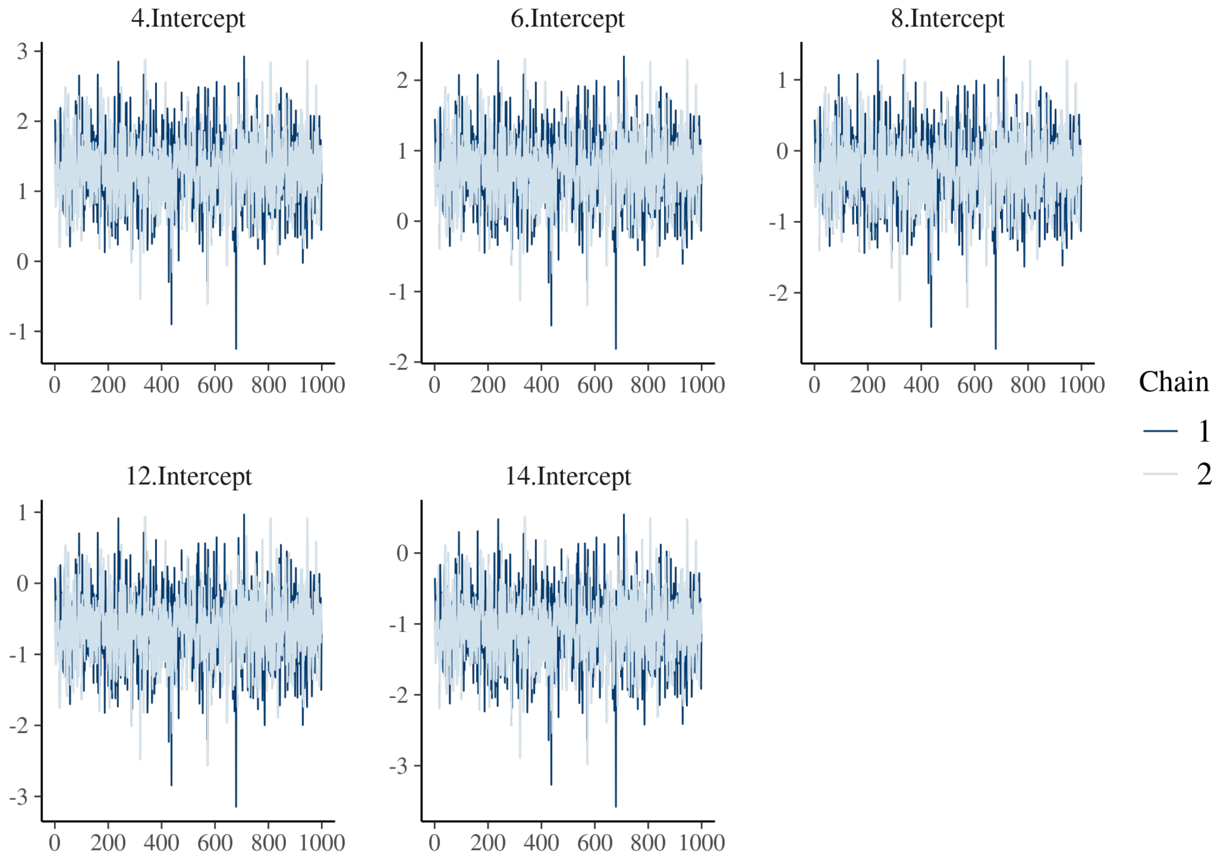

Figure S2.6: Trace of the temporal random effects

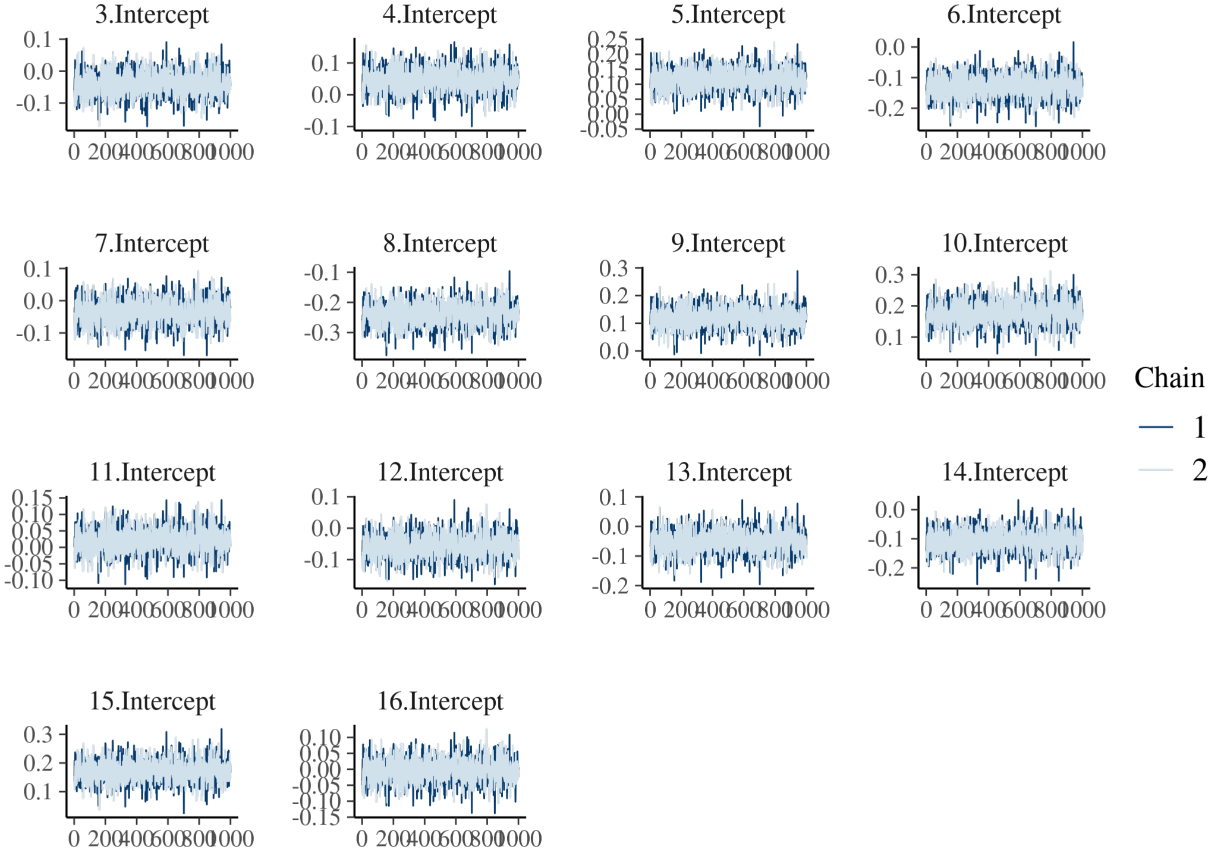

Figure S2.7: Trace of the genotype random effects

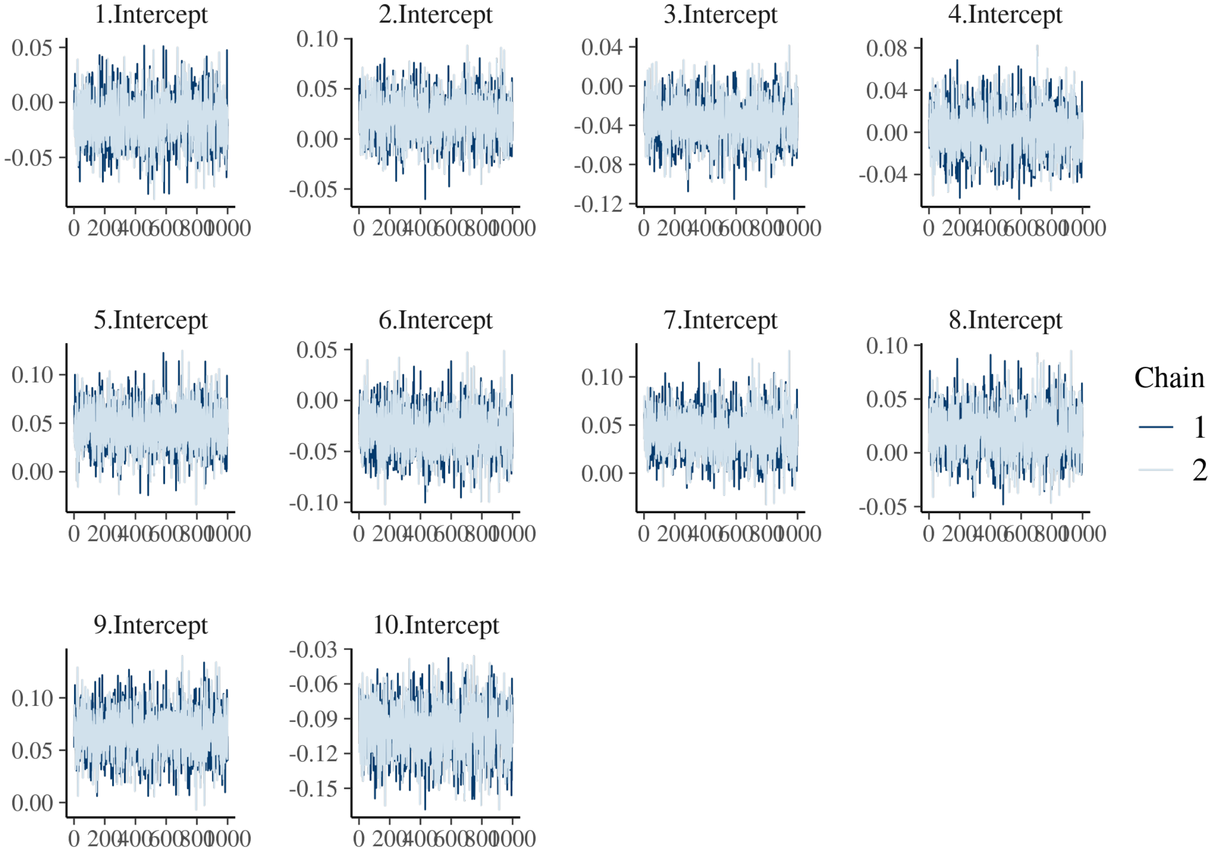

Figure S2.8: Trace of the spatial (block) random effects.

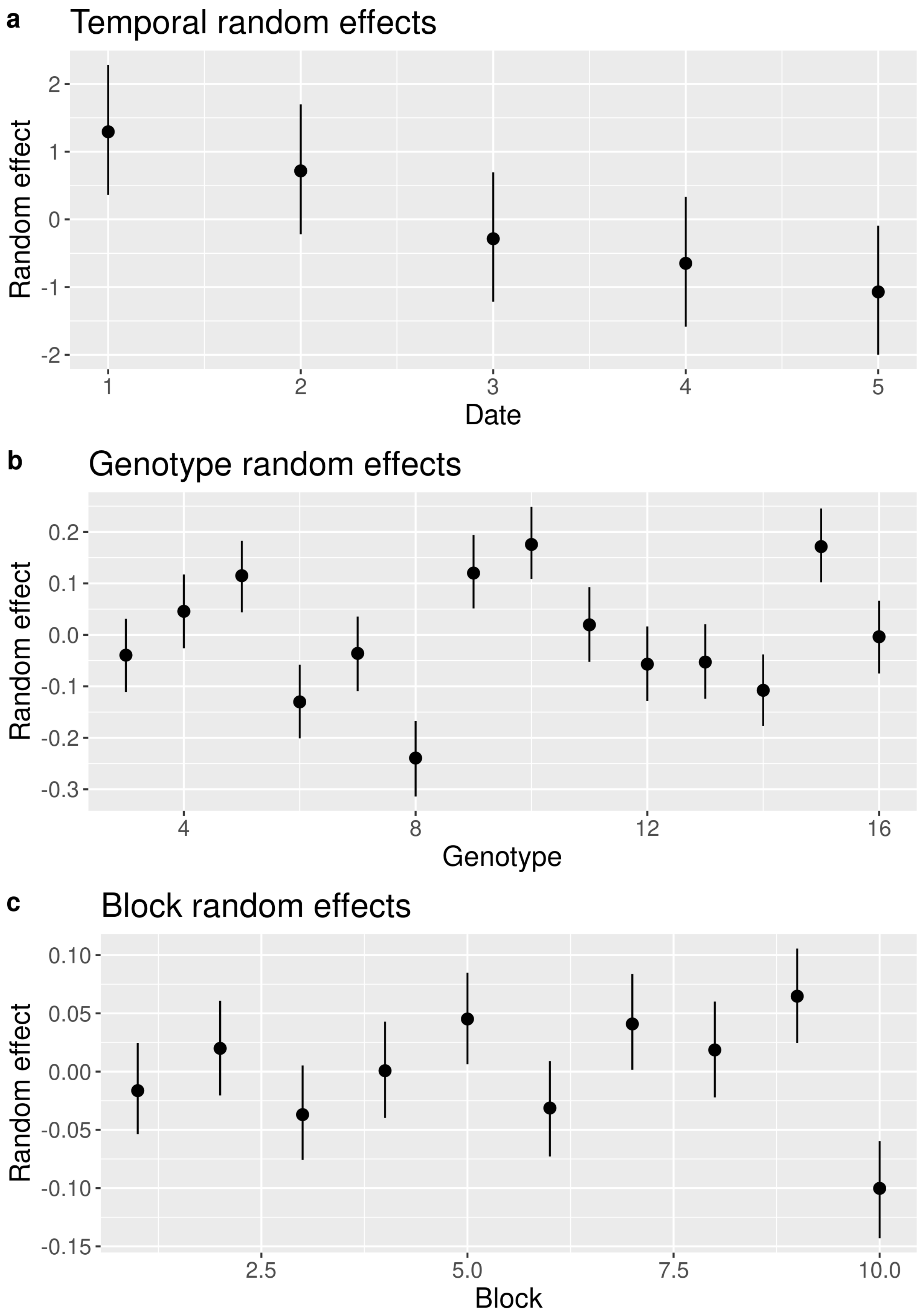

Figure S2.9: Mean values and 95% confidence interval of the temporal, genetic and spatial and random effects.

Figures S2.4 to S2.8 illustrate the convergence of the model.

Figure S2.9 shows mean values and 95% confidence intervals of random effects (except individual random effects which are too many), enabling to graphically look for a tendency.

### 3 Appendix 3: analysis of tropical forest inventory data

#### 3.1 Datasets

##### 3.1.1 Paracou

The Paracou forest is located in French Guiana. It is one of the best-studied lowland tropical forests in the Guiana Shield region. It belongs to the Caesalpiniaceae facies and has amongst the highest alpha-diversity in the Guiana shield with 150-200 tree species per hectare in inventories of trees with DBH $\geq$ 10 cm. The Guiana Shield is characterized by Pre-Cambrian granitic and metamorphic geological formations, highly eroded. It is associated with gently undulating landscapes and a very dense hydrographic system. Paracou forest lies in a hilly area, on a formation called "série Armina" characterized by schists and sandstones and locally crossed by veins of pegmatite, aplite and quartz. The topography of the site consists of small hills separated by narrow ($<$ 5 m wide) sandy waterbeds. The altitude varies from 5 to about 45 m above sea level Hérault and Piponiot (2018). The mean annual temperature is 26 °C and winds are generally weak. There is a well marked dry season (from mid-August to mid-November) and a long rain season with a short drier period between March and April (mean annual rainfall of 3,041 mm). Different study programs have been led at the Paracou site (https://paracou.cirad.fr/). Here, we used data from a disturbance experiment, where 12 square plots of 6.25 ha were exposed to four different logging intensities between 1986 and 1988, as well as four square plots of 6.25 ha which were set up for biodiversity monitoring in 1990-1992 (plots 1-15). Since then, cartesian coordinates, DBH, species identity and survival of each tree with a DBH $\geq$ 10 cm have been collected every one or two years, during the dry season (mid-August to mid-November). The studied 93.75-ha area harbors 91284 measured trees, among which 21688 belong to an undetermined species and others belong to 614 determined species, 221 genera and 64 families.

We removed all measures of the years before the perturbations were performed and the biodiversity plots were added (1985-1991), thus obtaining a dataset containing 91029 measured trees, among which 21481 belong to an undetermined species and others belong to 614 determined species, 221 genera and 64 families.

Figure S3.1 shows the location of the used trees in the Paracou site.

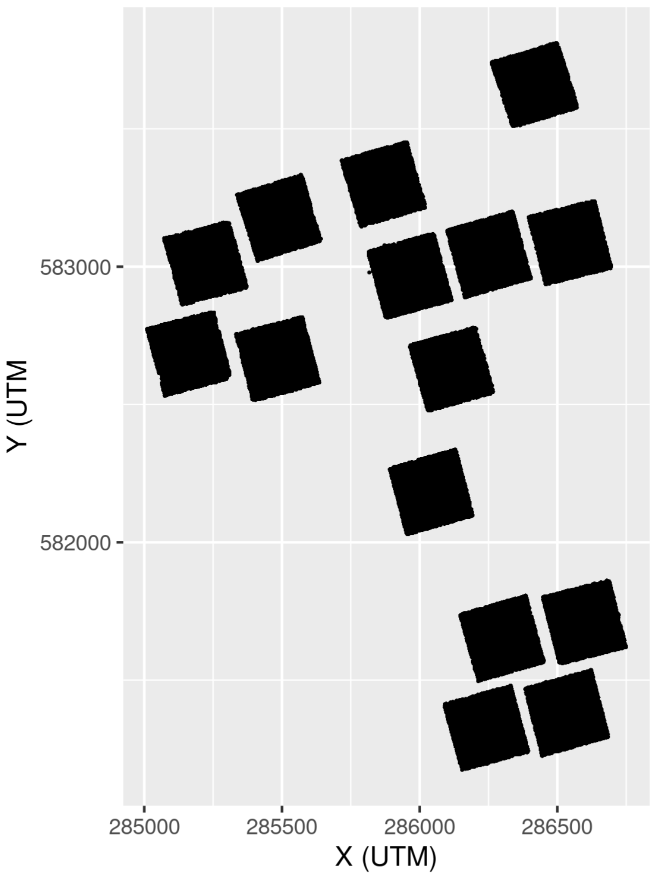

Figure S3.1: Plots at the Paracou site. Each point is a tree.

###### 3.1.2 Uppangala

The Uppangala Permanent Sample Plot (UPSP) is located in South-East Asia, in the Western Ghats of India, and was established in 1989 by the French Institute of Pondicherry in the Kadamakal Reserve Forest, in the Pushpagiri Wildlife Sanctuary, in Karnataka state, India (Pélissier et al. 2011). It is a low altitude (500-600 m) wet evergreen monsoon Dipterocarp forest (Bec et al. 2015). This forest is considered as one of the rare undisturbed tropical forests in the world (Pascal and Pelissier 1996). The studied area of 5.07 ha is quite steep, with a mean slope angle of about 30–35°. The plots consist in five North–South oriented transects that are 20-m wide, 180- to 370-m long, and 100-m apart center to center, in addition to three rectangular plots which overlap the transects. The transects gather data from 1990 to 2011 and the rectangular plots from 1993 to 2011. The trees with GBH (Girth at Breast Height) $\geq$ 30 cm (equivalent to ca. 9.5 cm DBH) were measured every 3 to 5 years. The original dataset contains measurements of 3870 trees of 102 species (including 2 morphospecies),78 genera and 32 families.

We removed the census dates which were not common for all plots (1990-1992).

Thus, we obtained a dataset containing 3870 trees of 102 species (including 2 morphospecies),78 genera and 32 families.

Figure S3.2 shows the location of the trees in the Uppangala site.

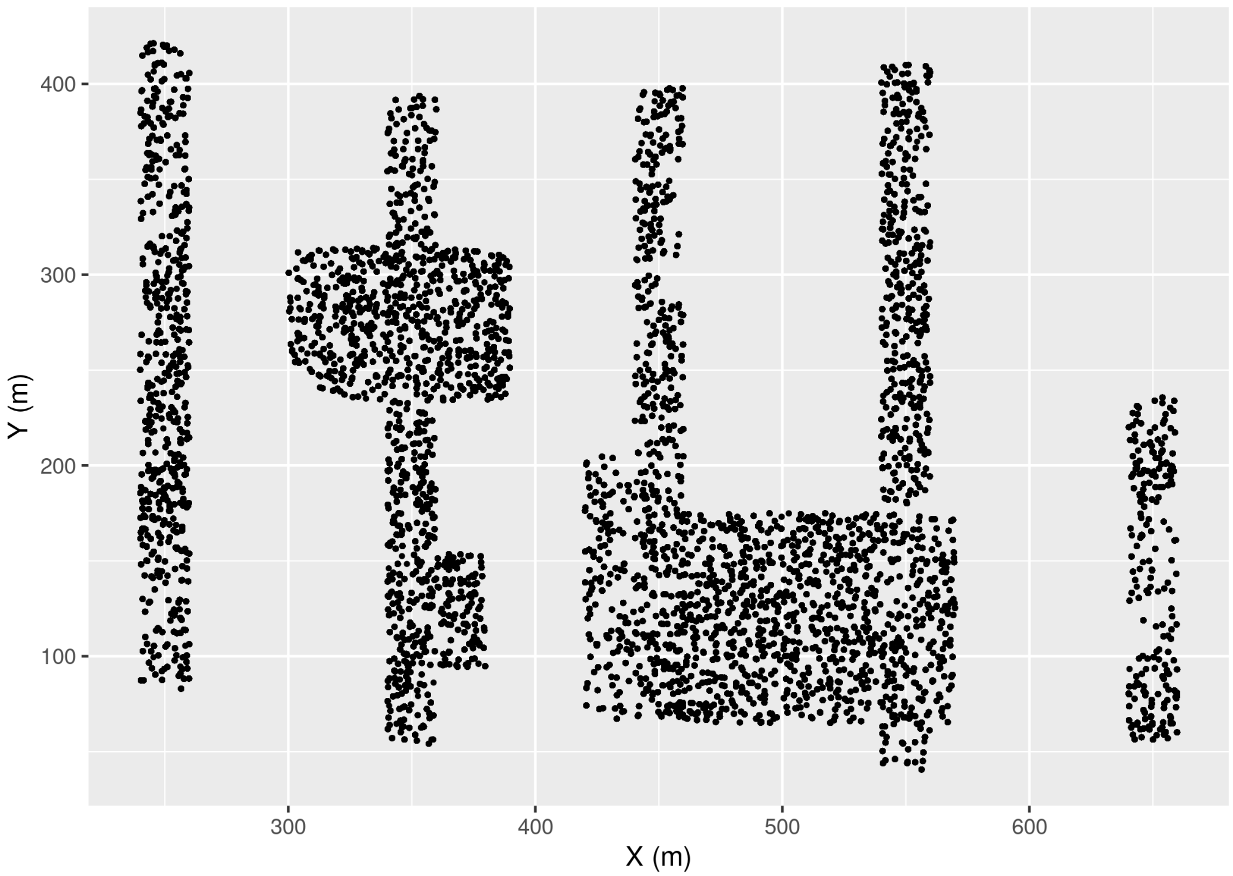

Figure S3.2: Plots at Uppangala site. Each point is a tree.

###### 3.1.3 BCI

The Barro Colorado Island site is located in central America, in Panama, covered by a lowland tropical moist forest. The zone became an island after a valley was flooded in order to build the Panama Canal, in 1913. It is nowadays considered as the most intensively studied tropical forest in the world. The studied site is a 50 ha plot (500 × 1,000 m). It has an elevation of 120 m and is quite flat (most slopes are gentler than 10°). Complete censuses of all trees with DBH $\geq$ 1 cm have been performed every 5 years since 1980. The dataset contains measurements of 423617 trees of 328 tree species, 195 genera and 63 families.

When only taking into account trees with DBH $\geq$ 10 cm for consistency with the other datasets, it contains measurements of 37224 trees of 255 tree species, 167 genera and 59 families. The dataset is available in (Condit et al. 2019).

Figure S3.3 shows the location of the selected trees in the BCI site.

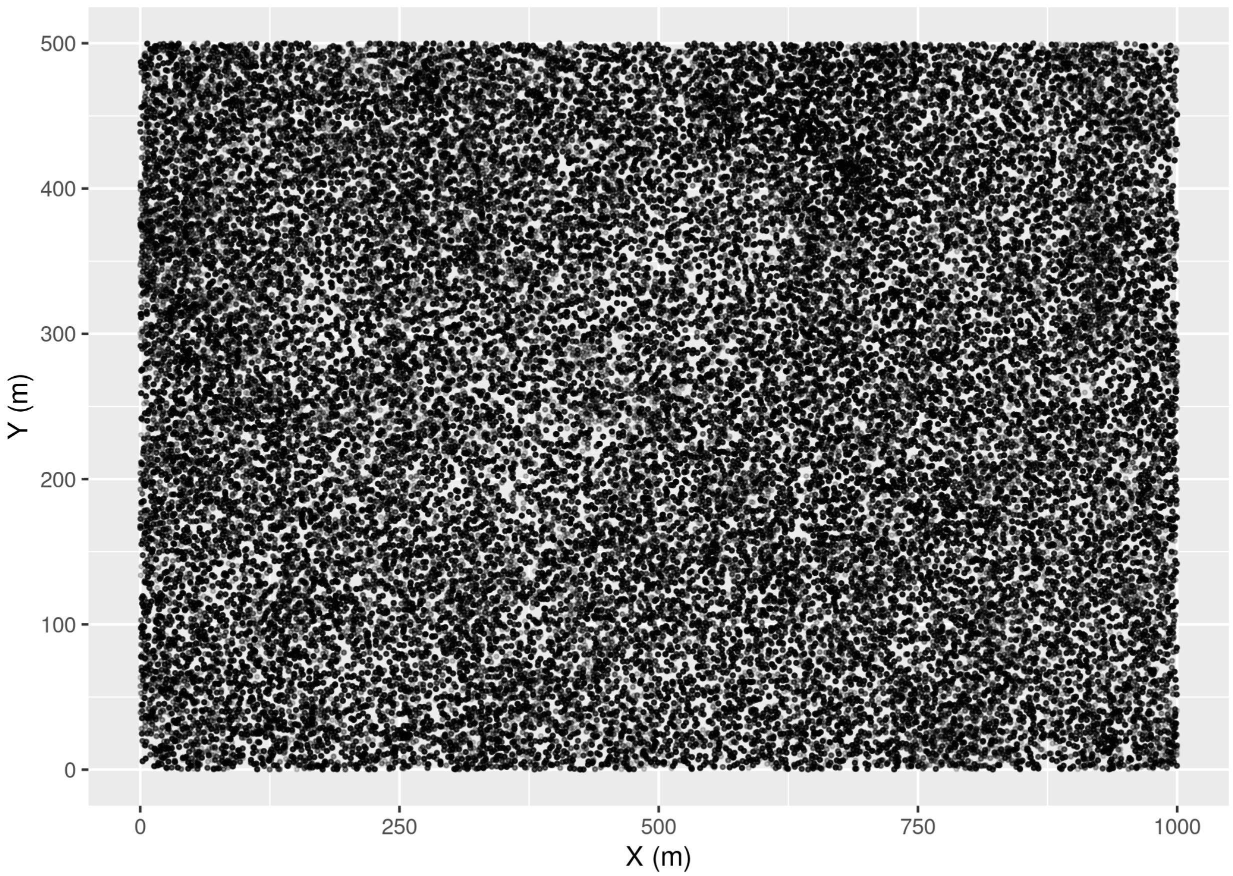

Figure S3.3: The 50 ha plot of the Barro Colorado Island site. Each point is a tree.

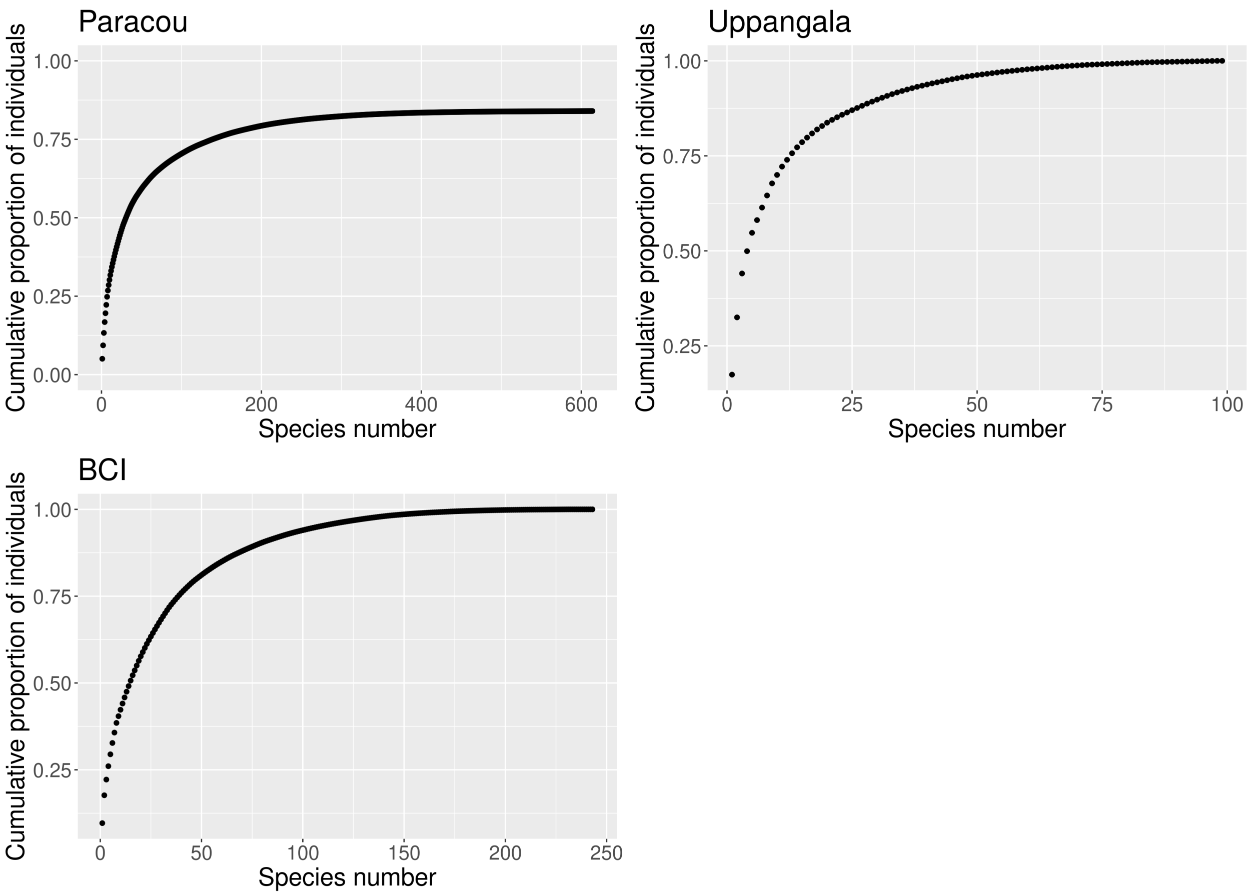

Figure S3.4: Abundance diagrams for the three study sites. For the Paracou site, only individuals that were identified to the species level are accounted for. The three communities include few dominant species and many rare species (long asymptote).

#### 3.2 Growth and mean growth estimation

For the Paracou site, after removing 105836 growth estimates for 12520 trees of indeterminate species, we obtained a dataset with 931389 growth estimates for 65769 trees of 614 species.

For the Uppangala site, we obtained a dataset with 57921 growth estimates for 3725 trees of 102 species.

For the BCI site, we obtained a dataset with 167618 growth estimates for 30386 trees of 244 species.

#### 3.3 Estimation of IV

We evaluated the convergence of the model using trace and density plots of the posterior estimates.

##### 3.3.1 Paracou

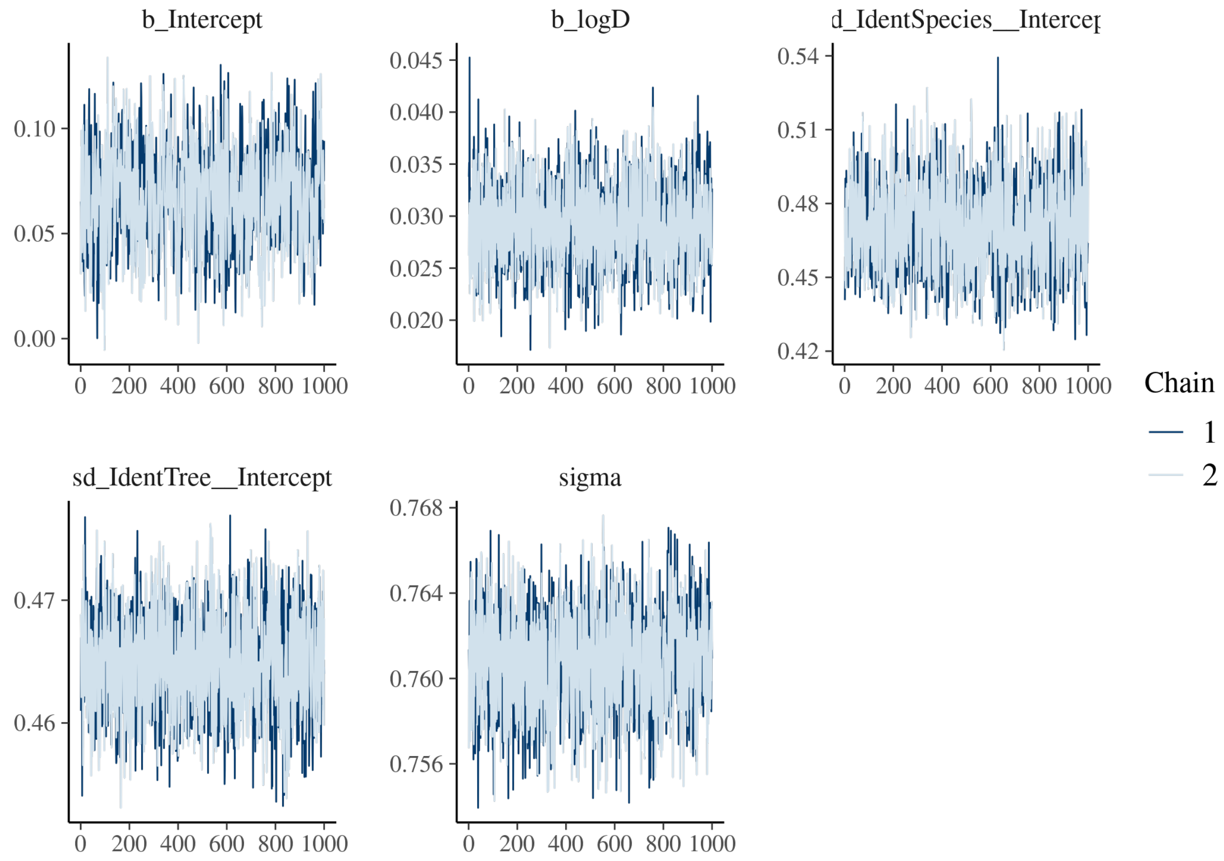

Figure S3.5: Trace of the posterior estimates for the Paracou site

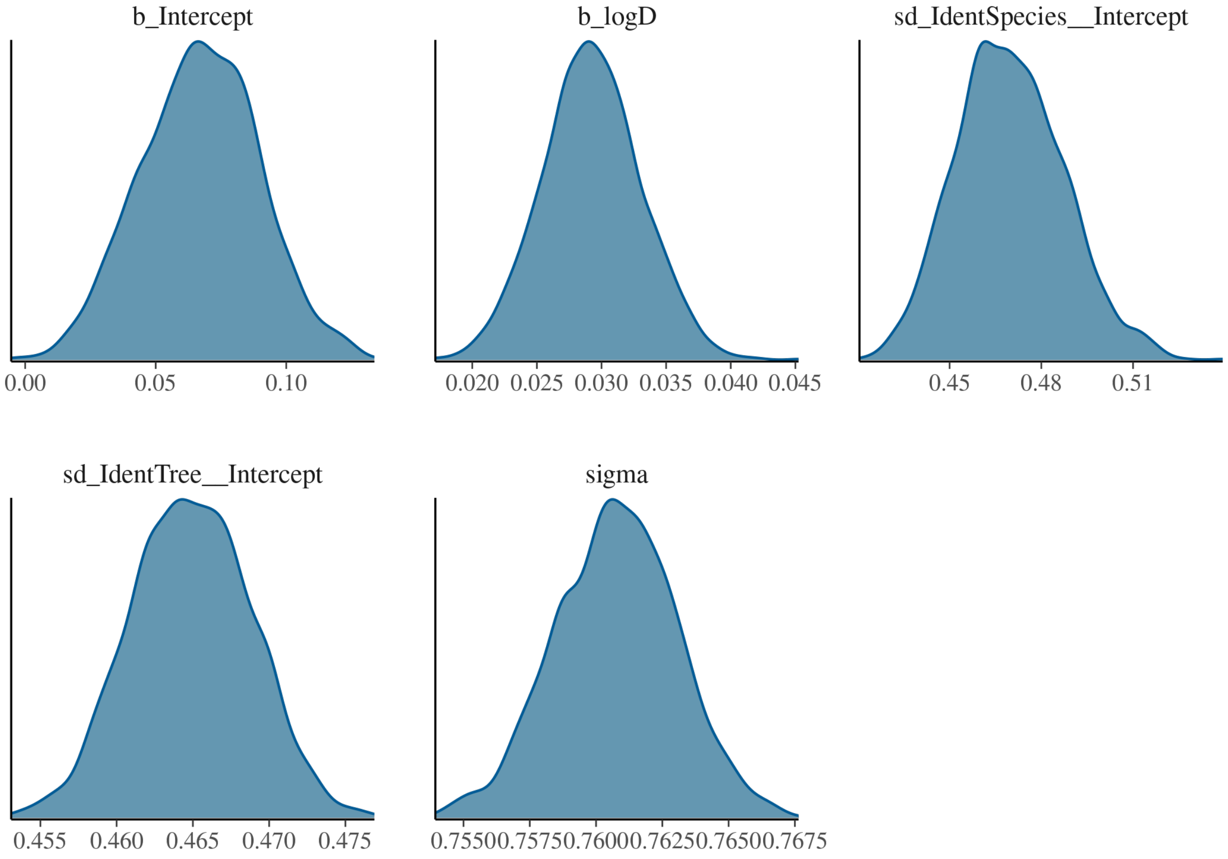

Figure S3.6: Density of the posterior estimates for the Paracou site

##### 3.3.2 Uppangala

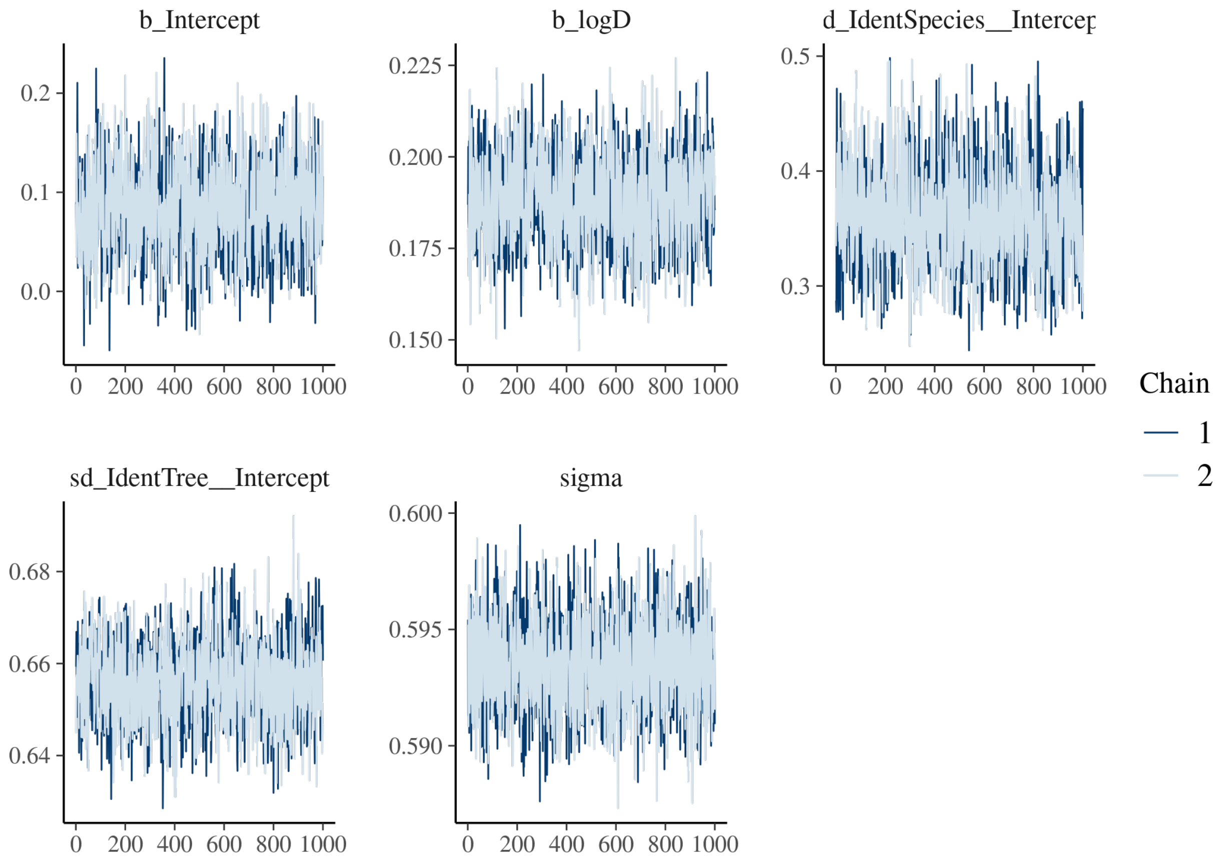

Figure S3.7: Trace of the posterior estimates for the Uppangala site

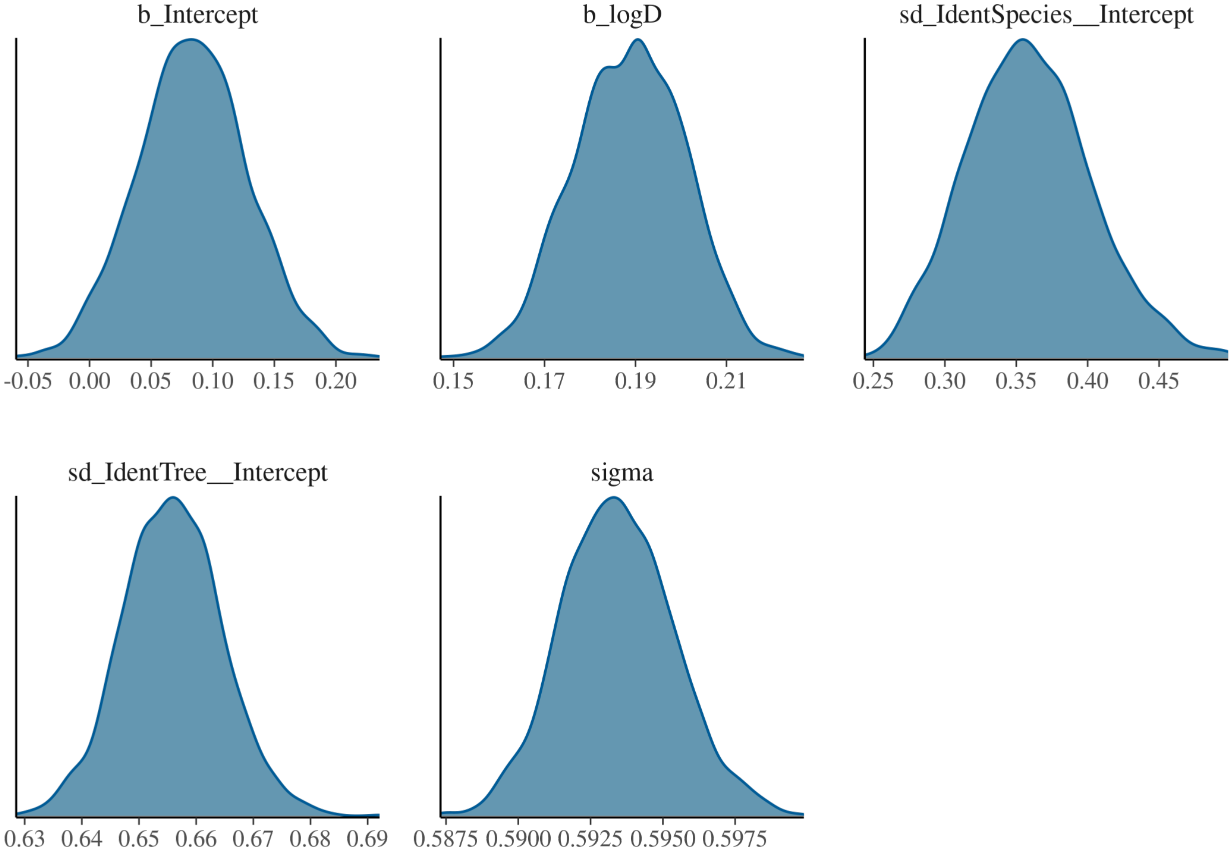

Figure S3.8: Density of the posterior estimates for the Uppangala site

##### 3.3.3 BCI

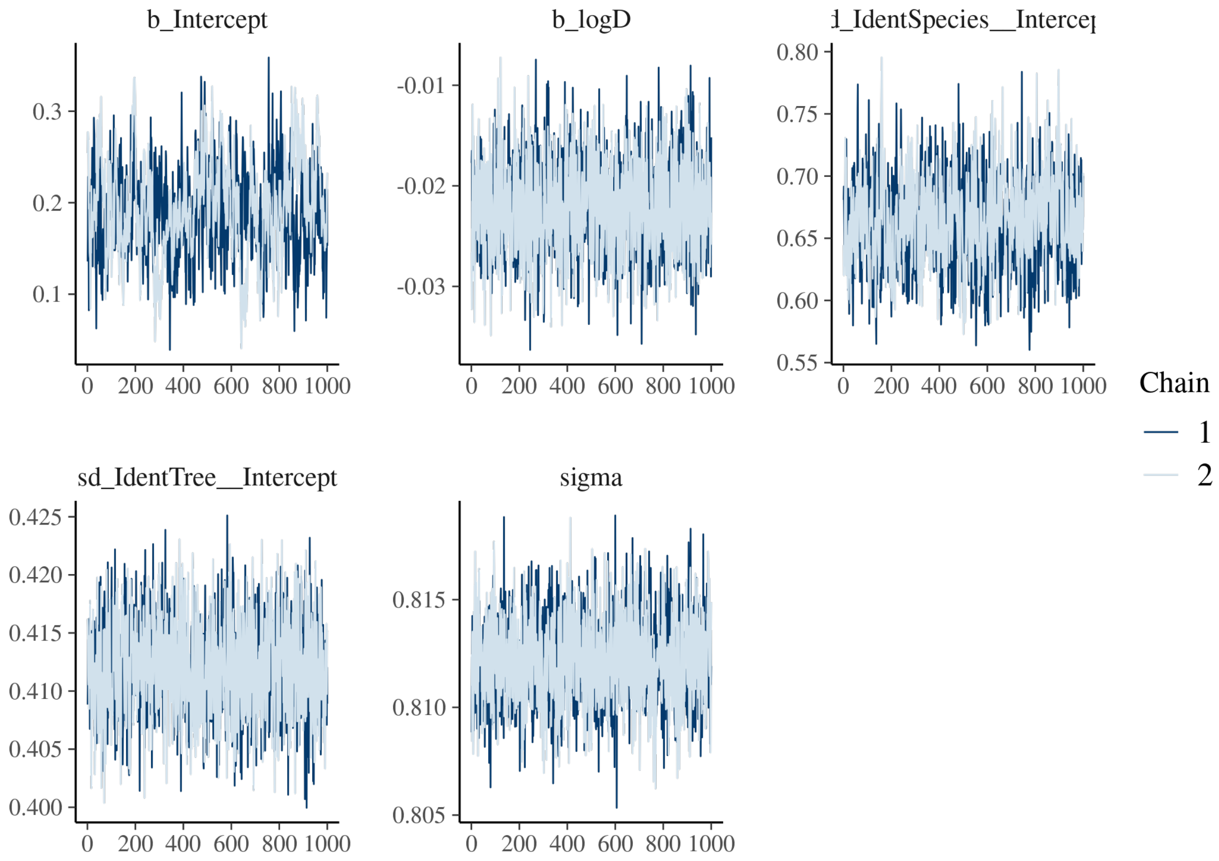

Figure S3.9: Trace of the posterior estimates for the BCI site

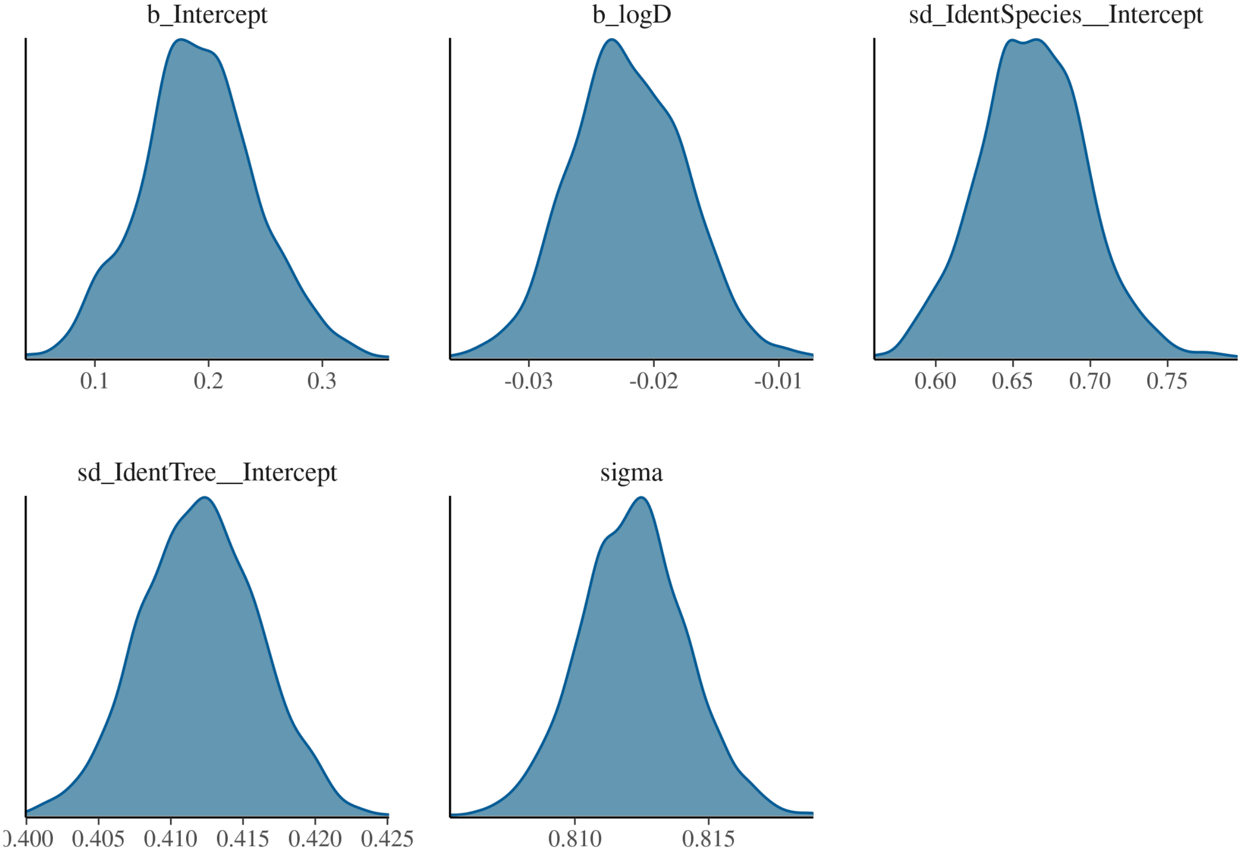

Figure S3.10: Density of the posterior estimates for the BCI site

#### 3.4 Spatial analyses

"Most abundant species" are species with more than 3,000 individuals.

#### 3.5 Spatial autocorrelation in undisturbed plots in the Paracou site

One of our main hypotheses was that since many environmental variables are spatially structured and that tree growth is largely influenced by environmental variables, tree growth should be spatially structured. Our analysis using Moran's I test showed that tree growth was indeed spatially structured at our study sites. The Paracou dataset offers the opportunity to test this hypothesis further. Indeed, some plots were disturbed in the early eighties, creating artificial gaps, whereas others were not disturbed. As the creation of gaps results in a strong spatial structure of the light available under the canopy, we hypothesized that growth should be less structured in plots that were not disturbed.

Considering the undisturbed plots only, the proportion of individuals of species with a significant Moran's I test was 55%, much lower than when including the disturbed plots (79%, see main text, Table 3; Table S3.1). This corroborates our hypothesis that the openness of the canopy triggered important spatial structure in disturbed plots due to light gaps, and as in these gaps, trees tend to grow faster, the spatial structure of growth is stronger.

Table S3.1: Spatial autocorrelation of the growth of conspecific individuals in the undisturbed plots of the Paracou site. Shown are the proportion of species, and of corresponding individuals, in %, for which individual growth was significantly spatially autocorrelated, or not significantly spatially autocorrelated. The spatial autocorrelation of individual growth was tested using Moran's I index.

| | Significant | Not significant |
|---|---|---|
| Proportion of species | 18.1 | 81.9 |
| Proportion of individuals | 54.7 | 45.3 |

##### 3.6 Spatial autocorrelation in BCI using smaller stems

In order to be able to compare all three datasets together, we removed all stems that had a DBH inferior to 10 cm in the BCI inventories, which include all stems with DBH $\geq$ 1 cm. We replicated our analysis on the spatial autocorrelation of tree growth in the complete dataset and found that the spatial structure in individual growth was even more significant.

##### 3.7 Variance comparison using a sample

Table S3.2: Comparison of local intra- and interspecific variability in individual growth for three tropical forest sites, when controlling for species abundance. See main text. Semivariances with heterospecifics were here computed by sampling a maximum of 10 individuals per species.

| | Intraspecific variability < interspecific variability (i) | Intraspecific variability ∼ interspecific variability (ii) | Intraspecific variability > interspecific variability (iii) |
|---|---|---|---|
| **Paracou** | | | |
| Proportion of species | 60.7 | 38.3 | 1 |
| Proportion of individuals | 86.7 | 12 | 1.26 |
| **Uppangala** | | | |
| Proportion of species | 42.2 | 55.6 | 2.22 |
| Proportion of individuals | 64.2 | 17.4 | 18.4 |
| **BCI** | | | |
| Proportion of species | 59.7 | 39 | 1.26 |
| Proportion of individuals | 88.7 | 10.7 | 0.516 |

To control for a potential effect of species abundance on the values of semivariances with heterospecifics, we replicated the analysis (see main text, Table 4) by computing the semivariances with heterospecifics by sampling a maximum of ten individuals per species. The results were qualitatively unchanged (Table S3.2).
